# Elucidating dynamic anaerobe metabolism with Live Cell HRMAS ^13^C NMR and genome-scale metabolic modeling

**DOI:** 10.1101/2022.06.13.495811

**Authors:** Aidan Pavao, Brintha Girinathan, Johann Peltier, Pamela Altamirano Silva, Bruno Dupuy, Isabella H. Muti, Craig Malloy, Leo L. Cheng, Lynn Bry

**Affiliations:** Massachusetts Host-Microbiome Center, Brigham and Women’s Hospital, Harvard Medical School; Boston, MA 02115, USA; Institut Pasteur, Université de Paris, UMR-CNRS6047; Laboratory of the Pathogenesis of Bacterial Anaerobes, 75015 Paris, France; Institute for Integrative Biology of the Cell (I2BC), CEA, CNRS, Université Paris-Saclay; 91198 Gif-sur-Yvette, France; Centro de Investigación en Enfermedades Tropicales, Facultad de Microbiología, Universidad de Costa Rica; San José, Costa Rica; Departments of Radiology and Pathology, Massachusetts General Hospital, Harvard Medical School; Boston, MA 02114, USA; Department of Radiology, The University of Texas Southwestern Medical Center; Dallas, TX 75390, USA; Clinical Microbiology Laboratory, Department of Pathology, Brigham and Women’s Hospital, Harvard Medical School; Boston, MA 02115, USA

## Abstract

Anaerobic microbial metabolism drives critical functions within global ecosystems, host-microbiota interactions, and industrial applications, yet remains ill-defined. Here we advance versatile approaches to elaborate dynamic metabolism in living cells of obligate anaerobes, using the pathogen *Clostridioides difficile*, an amino acid and carbohydrate-fermenting *Clostridia*. High-Resolution Magic Angle Spinning (HRMAS) Nuclear Magnetic Resonance (NMR) spectroscopy of *C. difficile* grown with uniformly labeled ^13^C substrates informed dynamic flux balance analysis (dFBA) of the pathogen’s genome-scale metabolism. Predictions identified metabolic integration of glycolytic and amino acid fermentation pathways at alanine’s biosynthesis, to support efficient energy generation, maintenance of redox balance, nitrogen handling, and biomass generation. Model predictions advanced an approach using the sensitivity of ^13^C NMR spectroscopy to simultaneously track cellular carbon and nitrogen flow, from [U-^13^C]glucose and [^15^N] leucine, confirming the formation of [^13^C,^15^N]alanine. We illustrate experimental and computational approaches to elaborate complex anaerobic metabolism for diverse applications.

## Main text

Obligate anaerobes comprise the majority of species in the mammalian gut microbiota and include pathogens such as *Clostridioides difficile*. Anaerobic bacteria also modulate nutrient flow across global ecosystems^1^ and perform industrial fermentations of economic importance^2^. However, the metabolic pathways and nutrient requirements of anaerobes often differ substantively from those of model, aerotolerant species such as *Escherichia coli* or *Bacillus subtilis*, leaving many aspects of their metabolism poorly defined. This poor characterization limits efforts to harness anaerobe metabolism in applications of clinical, industrial, or environmental improtance.

High-resolution magic angle spinning (HRMAS) NMR spectroscopy supports studies of real-time metabolism in living cells^3-5^, and is particularly suited to the study of anaerobes as the sealed rotor chamber can maintain an anaerobic environment^5^. HRMAS NMR rapidly rotates samples at a “magic angle” of 54.74° relative to the magnetic field during NMR spectrum acquisition, improving the sensitivity of signal detection in colloidal or semi-solid samples^6^. Detailed studies of metabolism can thus be achieved with a low input biomass of cells^5^. When coupled with cellular metabolism of uniformly carbon-13 (^13^C) labeled substrates, HRMAS NMR’s sensitivity enables definitive tracking of carbon flow through complex metabolic pathways.

Metabolic modeling systems link experimentally-obtained substrate and metabolite fluxes to cellular pathways and genes. By design, metabolic flux analyses (MFA) informed by ^13^C-NMR have remained limited to pathways carrying ^13^C flux^7-10^. In contrast, dynamic flux balance analysis (dFBA) simulates time-dependent recruitment of metabolic pathways on an organismal scale given a set of nutrient exchange constraints and a biological objective such as biomass or ATP production^11^. dFBA approaches estimate exchange fluxes from static measurements of the media composition over time^11,12^ for which repeat measurements by quantitative platforms, including GC/MS, have been used^13^. However, the use of NMR to constrain dFBA simulations has been limited by means to translate NMR signals into credible estimated concentrations, due to issues in NMR spectral resolution, signal-to-noise levels, and susceptibility to amplitude distortions from the Nuclear Overhauser Effect (NOE)^14^. With means to overcome these limitations, HRMAS NMR offers a promising approach to support dFBA given its non-destructive measurement of isotopic flux in minute quantities of living cells^5,15^.

*C. difficile*, the leading cause of hospital-acquired infections, colonizes gut environments through its metabolism of diverse carbon sources^16^. The pathogen releases toxins to obtain nutrients from damaged mucosa as its growth exceeds the carrying capacity of gut environments^17^. *C. difficile* ferments simple and complex carbohydrates, possesses multiple Stickland amino acid fermentation pathways^18-20^, and fixes carbon dioxide through Wood-Ljungdahl acetogenesis^21,22^, pathways shared with commensal and industrially-relevant *Clostridia*^13,21^. Defining how *C. difficile* recruits co-occurring fermentation pathways with systems supporting energy generation and growth has been challenging^23,24^ but offers opportunities to prevent and treat infections with approaches that need not rely solely on antibiotics.

We present a generalized approach using HRMAS ^13^C-NMR to constrain dFBA within genome-scale metabolic models, and define complex dynamics in anaerobe metabolism (Fig. 1a,b). Analyses link NMR signals with anaerobe pathways and genes, revealing metabolic integration points unique to amino acid fermenting *Clostridia*. Confirmation of findings further advanced a generalizable NMR approach using amplification of signal from weak NMR-active nuclei, such as ^15^N, through strong NMR-active nuclei, such as ^13^C, to track simultaneous carbon and nitrogen flow through cellular metabolism (Fig. 1c).

**Figure 1:**
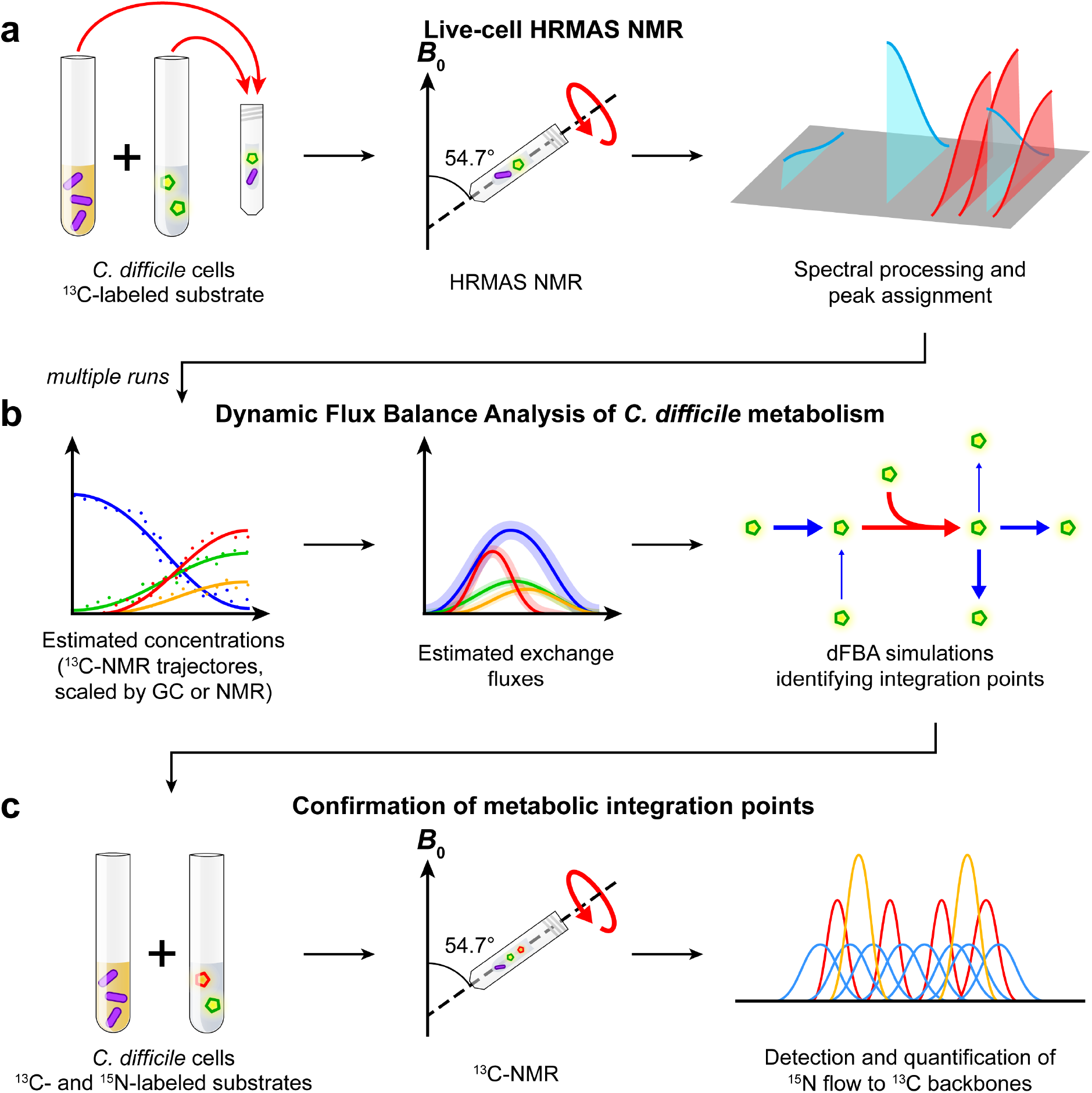
Framework for identifying metabolic integration points using HRMAS NMR with labeled substrates and dFBA. **(a)** HRMAS NMR resolves live-cell anaerobic metabolism of ^13^C-labeled substrates. A defined medium containing a ^13^C-labeled substrate is inoculated with *C. difficile* cells in an HRMAS rotor insert. Successive ^1^H- and ^13^C-NMR spectra of the growing cells are acquired throughout log-phase growth to monitor metabolism of the labeled substrate. NMR spectra are processed, and peaks are assigned to metabolites using ^1^H-^13^C HSQC spectra and reference data. **(b)** NMR signal trajectories inform dFBA simulations to identify metabolic integration points. Logistic curves for metabolites are fit to integrated ^13^C-signal trajectories and scaled to estimated concentrations using information from standard solutions measured by gas chromatography (GC) or NMR. Estimated metabolite exchange fluxes are derived from the logistic curves representing multiple NMR runs to constrain dFBA simulations. Metabolic integration points are identified where dFBA solutions predict substrates to exchange electrons or functional groups. **(c)** ^13^C-NMR of ^13^C- and ^15^N-labeled substrates confirms dFBA predictions of nutrient flow. *C. difficile* cells are grown in defined media containing ^13^C- and ^15^N-labeled substrates under NMR acquisition. ^15^N flow to ^13^C backbones is measured by quantifying the relative areas of split ^13^C-NMR subpeaks at the ^13^C-alanine alpha-carbon.

## Results

### Live-cell HRMAS NMR

To investigate the progression of glycolytic and amino acid fermentations in *C. difficile*, we measured HRMAS NMR time series of proton (^1^H) and ^13^C NMR spectra from living *C. difficile* cells. Cultures were grown in Modified Minimal Media (MMM)^25^ that replaced a given natural abundance carbon source with its uniformly-labeled carbon-13 isotopologue: L-[U-^13^C]proline, L-[U-^13^C]leucine, or [U-^13^C]glucose (Fig. 1a, Supplementary Table 1), nutrients known to drive rapid pathogen growth *in vivo*^24^.

As a Stickland fermenting *Clostridium*^*20*^, *C. difficile* prefers to metabolize simple sugars, such as glucose, and amino acids, such as proline and leucine, to sustain its metabolism and drive cellular growth^26,27^. *C. difficile’s* proline reductase reduces proline to 5-aminovalerate, coupling electron transfer to the bacterial Rnf complex, which supports ATP synthesis^18,28,29^. In media containing L-[U-^13^C]proline, 5-amino[^13^C]valerate production occurred over 5.9-16.5 hours, consuming 60% of proline (Fig. 2a,b).

**Figure 2:**
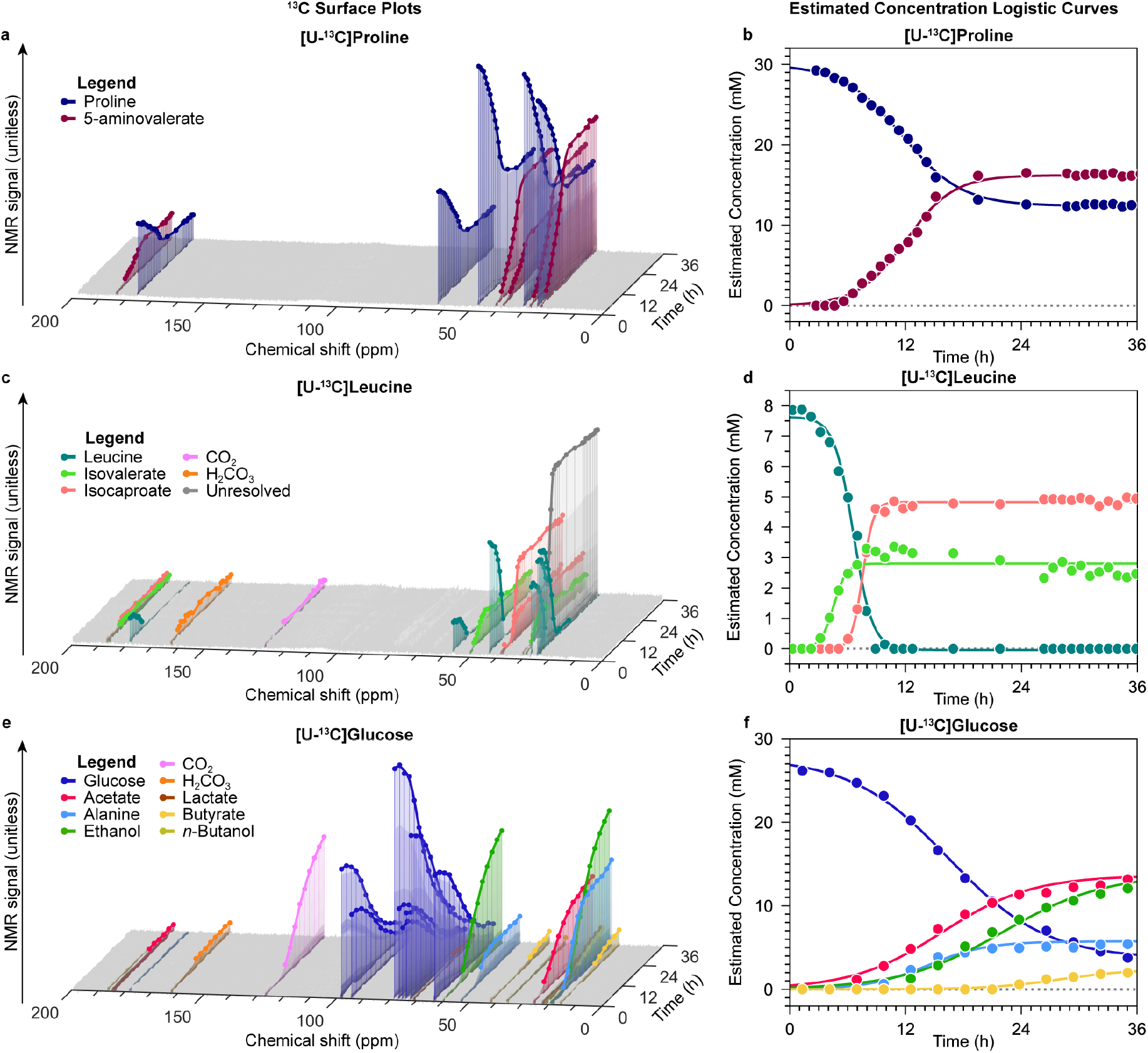
HRMAS ^13^C NMR of *C. difficile* grown with U-^13^C fermentable substrates. (**a**) HRMAS NMR stack plot of growth in MMM with 30mM [U-^13^C]proline. Legend shows color-coding of input proline (dark blue) and 5-aminovalerate (dark red). X-axis shows ^13^C NMR chemical shift (PPM), Y-axis: time (hours), and Z-axis: NMR signal (unitless). (**b**) Logistic plots depicting estimated concentration of [U-^13^C]proline and [U-^13^C]5-aminovalerate vs. time. (**c**) HRMAS NMR stack plot of growth in MMM with 7.6mM [U-^13^C]leucine. Legend shows carbons in leucine (teal) and detected metabolites; axes as in panel A. The 25 ppm peaks (gray) of isovalerate and isocaproate could not be resolved due to extensive overlap. (**d**) Logistic plots depicting estimated concentration of [U-^13^C]leucine and detected metabolites vs. time. (**e**) HRMAS NMR stack plot of growth in MMM with 27.5mM [U-^13^C]glucose. Legend shows color-coding of carbons in glucose (blue) and metabolites appearing over 36 hours of cellular metabolism; axes as in panel A. (**f**) Logistic plots depicting estimated concentration of [U-^13^C]glucose and detected metabolites vs. time.

*C. difficile* ferments L-leucine through oxidative and reductive pathways^19,27^. Oxidative fermentation of leucine to isovalerate and CO_2_ produces two reducing equivalents of ferredoxin and one equivalent of ATP, whereas the reductive leucine pathway yields isocaproate and consumes four net reducing equivalents^19,21,27^. Metabolism of L-[U-^13^C]leucine occurred within 13 hours, with levels of [^13^C]isovalerate and [^13^C]isocaproate rising over 2.7-8.7 hours and 7.4-13.0 hours, respectively (Fig. 2c,d).

HRMAS NMR of *C. difficile* grown with [U-^13^C]glucose identified [^13^C]acetate at 7 hours and [^13^C]alanine at 10 hours (Fig. 2e,f). Metabolic products of reductive glucose metabolism were detected by 13 hours with production of [^13^C]ethanol, followed by [^13^C]lactate at 21 hours, [^13^C]butyrate at 24 hours, and *n*-[^13^C]butanol at 35 hours (Fig. 2e,f).

### Estimating metabolic exchange fluxes from ^13^C-NMR trajectories

We employed three approaches to estimate flux constraints from the NMR time-series to use in dFBA supported by icdf843, an updated genome-scale metabolic model for *C. difficile*^30^. Estimates of changing metabolite concentrations first analyzed peak signals from the ^13^C-NMR spectra of the ^13^C-enriched components. The integrated NMR signal over time for each ^13^C-labeled compound was fit to a logistic curve, using the equation:

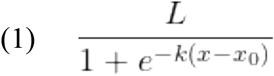

where *L* is the upper asymptote of the logistic curve, *k* is the growth rate of the curve, and *x*_0_ is the time of the inflection point of the sigmoidal curve (Supplementary Table 2). For the U-^13^C substrate, the constant *C* accounts for remaining substrate after 36 hours.

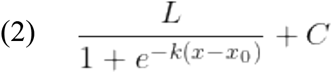

Curves were then differentiated to obtain estimated changes in the input ^13^C substrate or ^13^C metabolites:

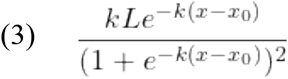

While the direct relationship between relative proton abundance and peak area in the ^1^H-NMR spectrum can be leveraged to estimate the relative abundance of protons at specific molecular contexts in a sample, an equivalent relationship between ^13^C abundance and peak area does not exist in ^13^C-NMR due to molecular context effects, including Nuclear Overhauser Effect (NOE) cross-relaxation between adjacent protons and the ^13^C nuclei^14^. This property of ^13^C-NMR prevents concentration estimates using relative signal intensities alone.

To overcome limitations in ^13^C-NMR for estimating metabolite concentrations to constrain exchange fluxes, the following approaches were used. First, the ^13^C substrate’s logistic derivative equation (3) was scaled by its input concentration in MMM, resulting in an equation for estimated uptake flux of the substrate. We next evaluated the metabolites produced during a labeled substrate’s fermentation and their predicted origin(s) from single or multiple metabolic pathways. If pathway associations were unknown or incompletely defined, the more complex case of multiple pathway origins was used.

In the case of a single pathway relationship between the ^13^C substrate and product, such as with *C. difficile’s* fermentation of proline to 5-aminovalerate^31^, the product formed was scaled relative to the substrate consumed and its flux calculated accordingly.

In contrast, leucine fermentation proceeds to two metabolites, isovalerate and isocaproate, that each originate from unique metabolic pathways in *C. difficile*^19,20^. As the inconsistency of NMR sensitivity among carbon contexts limits determination of the ratio of these metabolites by ^13^C-NMR, an expected ratio of isovalerate to isocaproate was determined by gas chromatography (GC, Supplementary Table 3), an alternate and quantitative method^24^, to scale the ^13^C-isovalerate and ^13^C-isocaproate trajectories from the input ^13^C-leucine concentration.

In more complex cases where fermentation products are not exclusive to the ^13^C substrate, a method able to discriminate ^13^C from ^12^C-produced metabolites is required. To estimate metabolite concentrations from [U-^13^C] glucose, standard solutions containing ^13^C products of glucose fermentation (Supplementary Table 4) were measured by ^13^C-NMR to quantify the relative ^13^C-NMR sensitivity of each metabolite with respect to glucose (Extended Data Fig. 1). The logistic derivative equations (3) were then scaled by the input glucose concentration and each product’s experimentally determined ^13^C-NMR signal enhancement factor, yielding estimated exchange fluxes over time to serve as constraints for dFBA (Fig. 2b,d,f; Supplementary Table 5; see Methods, “Estimation of exchange fluxes for dFBA”).

### Dynamic Flux Balance Analysis of *C. difficile* metabolism

Each of these methods to estimate metabolite concentrations from ^13^C-NMR signals informed dFBA to link HRMAS ^13^C NMR signals to the supporting metabolic pathways, genes, and their recruitment over time (Fig. 1b). Model predictions, with ATP hydrolysis as the biological objective (Fig. 3a), defined estimated metabolite flux at each timepoint and over the time series (Supplementary Table 6).

**Figure 3:**
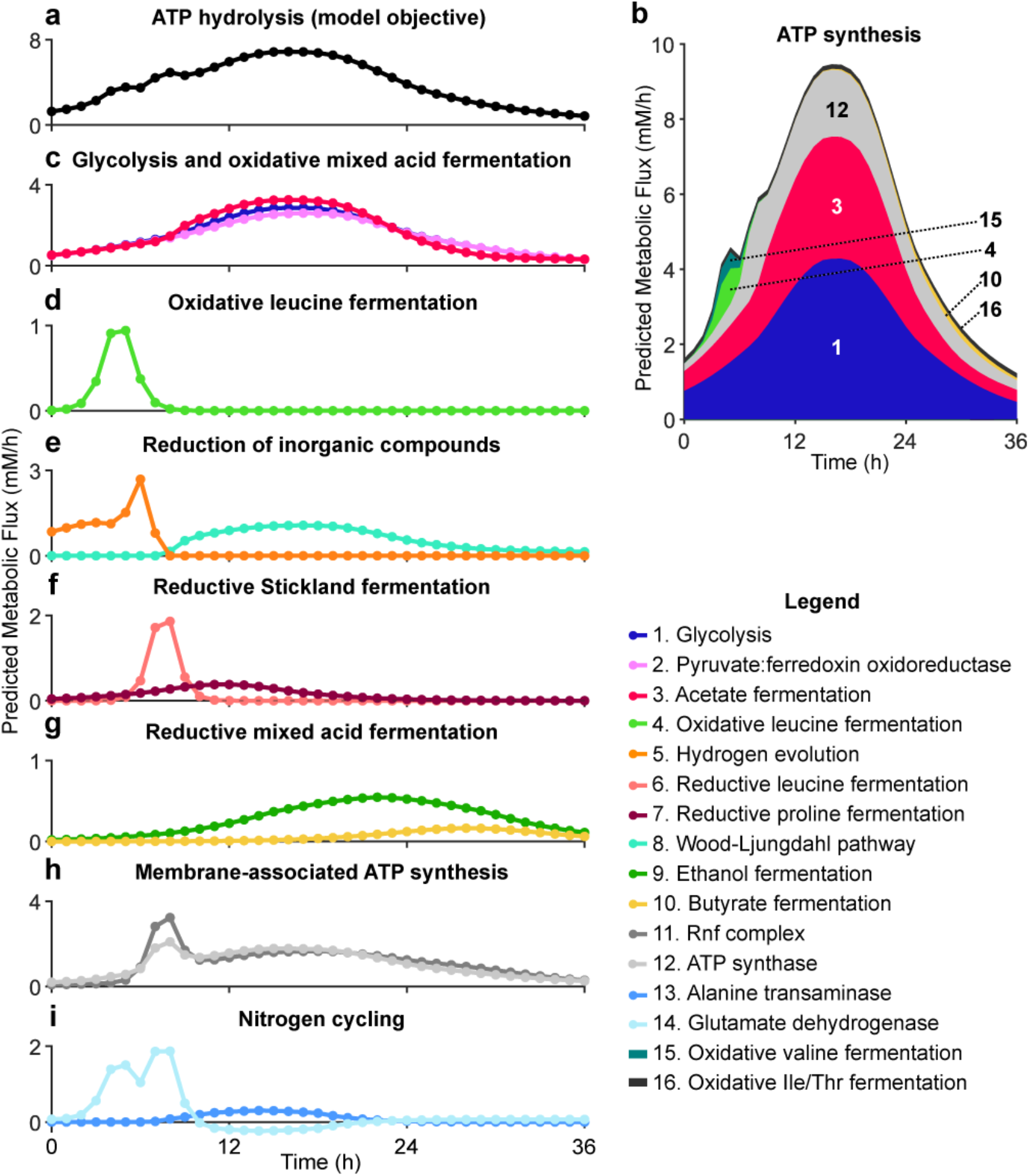
dFBA predicted reaction fluxes and production of key metabolic intermediates linking glycolysis and Stickland metabolism. (**a-i**) dFBA predicted reaction fluxes over 36h of metabolism (X-axis). Y-axis shows inferred flux in mM/h. (**a**) ATP hydrolysis, the model objective. (**b**) Inferred flux of reactions contributing to the production of ATP vs. time, including glycolysis represented by phosphoglycerate kinase and pyruvate kinase, acetate fermentation represented by acetate kinase, gradient-driven ATP synthase, oxidative leucine fermentation represented by isovalerate kinase, oxidative valine fermentation represented by isobutyrate kinase, butyrate fermentation represented by butyrate kinase, and other oxidative Stickland fermentations represented by propionate kinase and 2-methylbutyrate kinase. (**c**) Estimated glycolytic flux represented by Enolase and oxidative mixed acid fermentation represented by pyruvate:ferredoxin oxidoreductase (PFOR) and acetate kinase, (**d**) oxidative leucine fermentation represented by 2-oxoisocaproate dehydrogenase, (**e**) inorganic reductions represented by hydrogen evolution and the Wood-Ljungdahl acetyl-CoA synthase, (**f**) reductive Stickland fermentation represented by 2-isocaprenoyl-CoA dehydrogenase and proline reductase, (**g**) reductive mixed-acid fermentation represented by ethanol dehydrogenase and butyrate kinase, (**h**) membrane-associated ATP synthesis represented by ATP synthase and the gradient-producing Rnf complex, and (**i**) nitrogen cycling via alanine transaminase and glutamate dehydrogenase. Positive glutamate dehydrogenase flux depicts the forward reaction oxidizing glutamate to 2-oxoglutarate; negative flux, below the X-axis, depicts the reverse reaction.

Analyses predicted flux through oxidative reactions driving ATP synthesis (Fig. 3b), with co-occurring reductive reactions accepting electrons to sustain oxidative flux. Model simulations predicted oxidative reactions among glycolysis (Fig. 3b,c, Fig. 4, reaction 1) and mixed acid fermentation (Fig. 3b,c, Fig. 4, reactions 2 and 3) to drive ATP production over 36 hours of metabolism, with 1.5% of ATP synthesis from oxidative leucine fermentation occurring during the first 8 hours of metabolism (Fig. 3b,d, Fig. 4, reaction 4; Supplementary Table 6).

**Figure 4:**
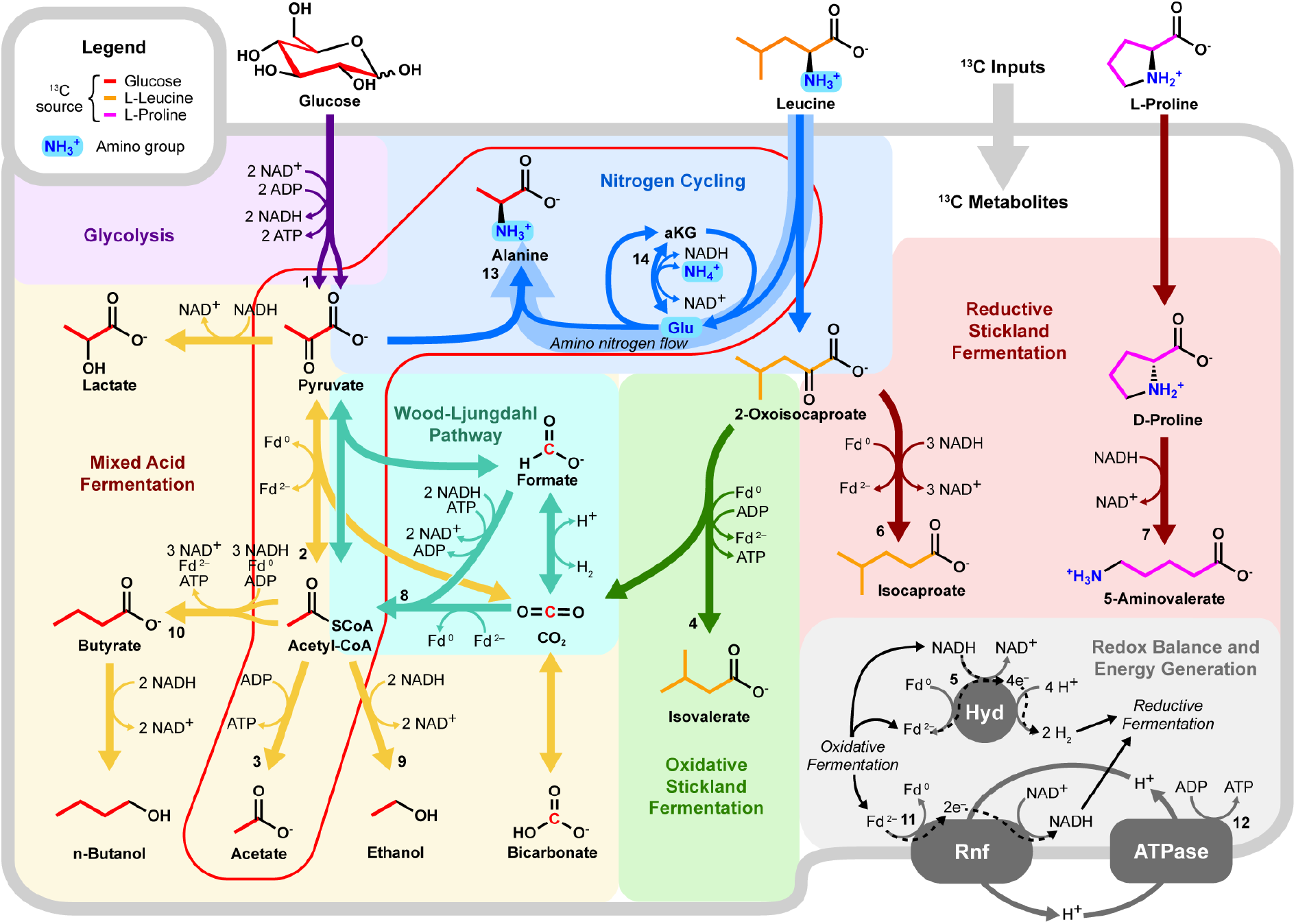
HRMAS NMR and dFBA-informed metabolic map for *C. difficile*. Diagram shows an updated metabolic map for *C. difficile* highlighting alanine’s role in nitrogen cycling (blue) and integration of glycolytic and mixed acid fermentations (yellow) with Stickland oxidative (green) and reductive (red) reactions that release abundant amino nitrogen. The map also includes the Wood-Ljungdahl pathway (teal) and iron hydrogenase (“Hyd”), predicted by dFBA to serve as key electron sinks, and the Rnf-ATPase system responsible for ATP-producing electron transfer between oxidative and reductive fermentations (gray). Numbered reactions correspond to the reaction legend in Figure 3 and Supplementary Table 8. Red-lined box indicates central reactions linking pyruvate, acetate, and alanine across glycolytic and Stickland fermentations.

Solutions predicted progressive stages of reductive metabolism, first with the evolution of molecular hydrogen during the first 7 hours of metabolism (Fig. 3e, Fig. 4, reaction 5), then by the Stickland reduction of leucine and proline (Fig. 3f, Fig. 4, reactions 6 and 7). After the rapid depletion of Stickland acceptors, dFBA solutions predicted the Wood-Ljungdahl pathway (Fig. 3e, Fig. 4, reaction 8) and ethanol fermentation (Fig. 3g, Fig. 4, reaction 9) as the primary electron sinks beginning at 10 hours. By 24 hours, late-stage reductive metabolism shifted to reactions producing butyrate and *n-*butanol, contributing 1.2% of overall ATP synthesis via substrate-level phosphorylation (Fig. 3b,g, Fig. 4, reaction 10; Supplementary Table 6). dFBA simulations predicted that the Rnf complex (Fig. 3h, Fig. 4, reaction 11, “Redox Balance and Energy Generation”) harnessed the energy cascade between oxidative and reductive processes to generate proton gradients and ATP production through *C. difficile’s* F-type ATP synthase (Fig. 3b,h, Fig. 4, reaction 12).

dFBA solutions predicted alanine’s production (Fig. 3i, Fig. 4, reaction 13) as a central integration point to support concomitant glycolytic and Stickland amino acid fermentations in *C. difficile* for energy generation, nitrogen handling, and cellular growth (Fig. 4, red-lined box). While model solutions predicted net oxidative deamination of amino acids during the first 9 hours of metabolism (Fig. 3i, Fig. 4, reaction 14), leucine-origin nitrogen, recycled onto glutamate between 10 and 21 hours, was predicted to supply up to 70% of amino group nitrogen in alanine biosynthesis (Fig. 4, “Amino nitrogen flow”; Supplementary Table 6), identifying alanine as a small molecule link between glycolysis and Stickland amino acid fermentations.

### Confirmation of metabolic integration between glycolytic and Stickland metabolism

To confirm the model predictions of amino nitrogen flow from Stickland leucine fermentation to ^13^C pyruvate, we developed an NMR approach to simultaneously track cellular carbon and nitrogen flow. Tracking of less sensitive NMR-active nuclei, such as nitrogen-15 (^15^N), through cellular metabolism has been more challenging than that of more sensitive NMR-active nuclei such as ^13^C and ^1^H. Nitrogen-15 (^15^N) produces 15-fold less signal than ^13^C^32^. However, NMR J-coupling between ^13^C and covalently bound ^15^N induces predictable patterns of nuclear spin-spin splitting in ^13^C signals^33,34^, enabling detection of the less sensitive ^15^N nucleus in the more sensitive ^13^C NMR spectrum.

Leveraging this property of NMR physics, we tracked simultaneous flow of ^13^C backbones from [U-^13^C]glucose and ^15^N amino groups from [^15^N]leucine to alanine (Fig. 1c, Fig. 4, “Nitrogen cycling”). Nitrogen-15 (^15^N, 0.4% natural abundance^35^) has a nuclear spin of +1/2 and splits a bonded ^13^C signal into two equivalent peaks in ^13^C NMR due to the J-couplings between ^15^N and ^13^C^33,34^. The increased mass of the ^15^N nucleus also causes an additional upfield shift of the bonded ^13^C peak, known as the isotope effect^36^. These combined effects enable discrimination of [^13^C,^15^N]alanine from [^13^C,^14^N]alanine by ^13^C NMR.

To confirm the feasibility of this approach, we first evaluated ^13^C-induced splitting of acetate’s peaks in the ^1^H spectrum of cells grown with [U-^13^C]glucose. Signal from the methyl hydrogens of [^13^C_2_]acetate was split into double-doublet peaks, indicating well-resolved ^1^H spin coupling with the ^13^C nucleus in the methyl group (*J* ∼ 34 Hz), and long-range coupling with the ^13^C nucleus in the carboxyl group (*J* ∼ 53 Hz; Extended Data Fig. 1b, Methods). Moreover, the amplification of ^13^C signal in the ^1^H spectrum showed 30 times higher signal-to-noise ratio (S/N) than in the ^13^C spectrum alone (Extended Data Fig. 2b, Methods), demonstrating that spin-spin splitting in a more sensitive NMR spectrum (^1^H) offers enhanced detection of comparatively less sensitive NMR nuclei (^13^C).

We utilized this advantage of NMR physics to amplify less sensitive ^15^N signal in the ^13^C NMR spectrum by detecting ^15^N-^13^C bonds occurring in alanine. HRMAS ^13^C NMR time series of *C. difficile* grown with [U-^13^C]glucose and natural abundance leucine revealed [2,3-^13^C]alanine and [U-^13^C]alanine in a 1:1 ratio (Fig. 5a,b; Extended Data Fig. 3; Supplementary Table 7), indicating substantial assimilation of ^12^CO_2_ with [U-^13^C]acetate, an activity reported to occur via pyruvate:ferredoxin oxidoreductase in many species of *Clostridia*^37^. *C. difficile* grown in the presence of [U-^13^C]glucose and [^15^N]leucine showed ^15^N-induced splitting (*J* ∼ 5.6 Hz) and isotope effect shifting (*δ* ∼ 0.025 ppm) of the ^13^C peaks associated with alanine’s alpha carbon and mixed populations of [^15^N]alanine and [^14^N]alanine (Figs. 5c,d; Extended Data Fig. 4). Though [^15^N]leucine represented only 33% of amino-group nitrogen in the starting media, 57±4% of [^13^C]alanine carried the ^15^N isotope (Fig. 5e), confirming enriched transfer of the ^15^N amino group from fermented [^15^N]leucine to [^13^C]alanine (Fisher’s Exact Test, p=0.001).

**Figure 5:**
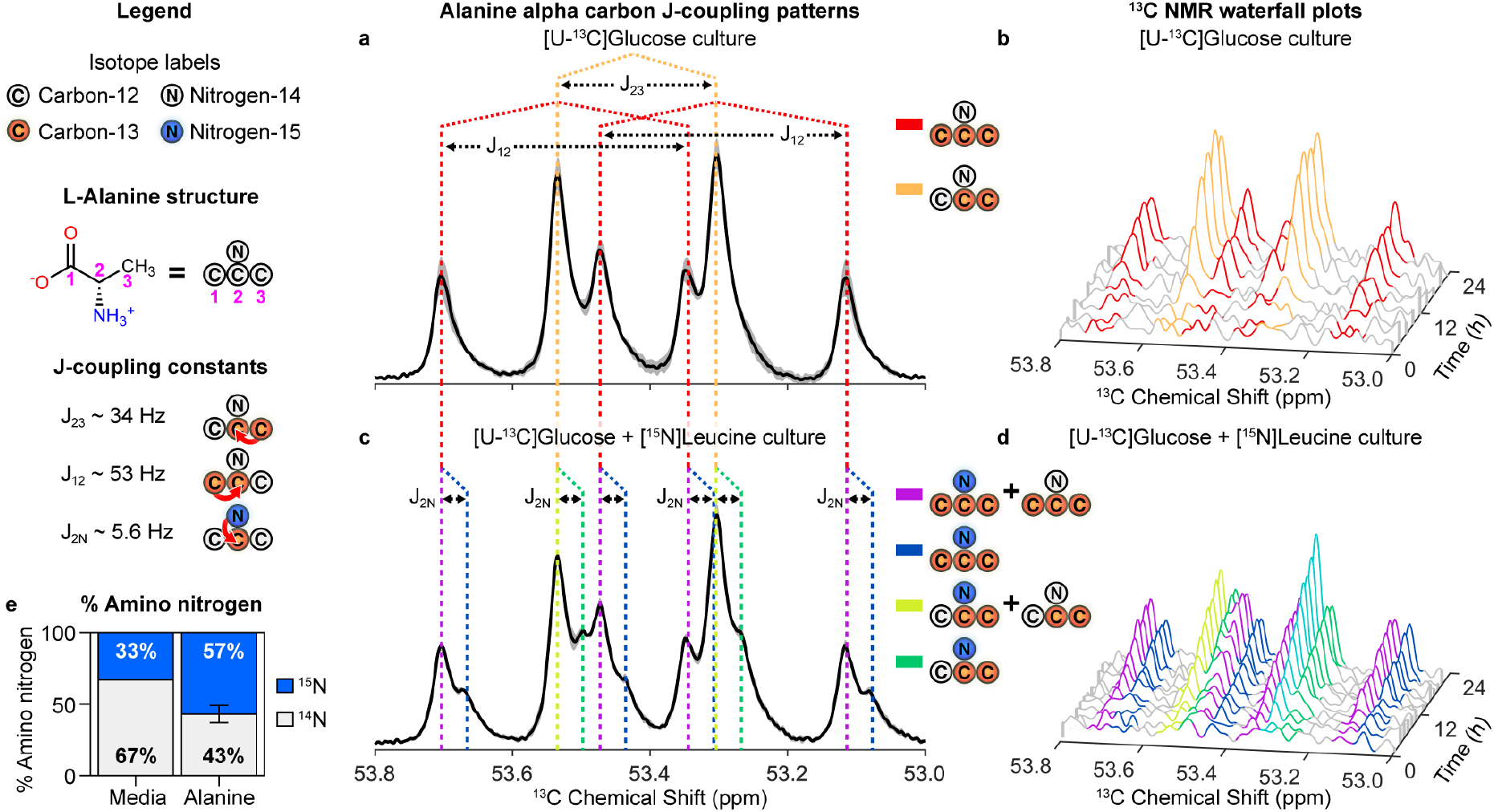
^13^C NMR detects ^15^N amino nitrogen flow from Stickland fermented leucine to glycolytic ^13^C backbones in alanine. (**a**) ^13^C NMR spectra for alanine’s alpha carbon (53.0-53.8 ppm) after growth in MMM media with 27.5mM [U-^13^C]glucose and 7.6mM natural abundance leucine for 36 hours. Red-dotted lines indicate J-coupled peaks associated with [U-^13^C]alanine. Orange-dotted lines indicate J-coupled peaks associated with [2,3-^13^C]alanine resulting from ^12^CO_2_ assimilation with [U-^13^C]acetate, via pyruvate:ferredoxin oxidoreductase (PFOR). (**b**) Time series of J-coupled peaks at alanine’s alpha carbon over 24h of metabolism, color coded as shown in panel (B). (**c**) ^13^C NMR spectra after growth in 27.5mM [U-^13^C]glucose and 7.6mM [^15^N]leucine showing J-coupled split peaks from [U-^13^C,^15^N] alanine (blue) or [2,3-^13^C^,15^N] alanine (green). Purple lines show peaks with a mixture of [U-^13^C,^15^N]alanine and [U-^13^C, ^14^N]alanine; chartreuse lines show peaks with [2,3-^13^C^,15^N]alanine and [2,3-^13^C^, 14^N]alanine. (**d**) Time-series of J-coupled peaks, color coded as shown in panel (C). *Asterix over aqua-colored peak indicates combined [2,3-^13^C,^14^N]alanine, [2,3-^13^C,^15^N]alanine, and [U-^13^C, ^15^N]alanine. (**e**) Table shows percentages of ^14^N and ^15^N amino nitrogen in starting media (Supplementary Table 7) with [U-^13^C]glucose and [^15^N]leucine, and resulting integrated ^14^N-^13^C and ^15^N-^13^C percentages (±SEM) on alanine after 36h of growth; Fisher’s exact test p=0.001.

## Discussion

We illustrate advancements in HRMAS NMR and genome-scale metabolic modeling to track single carbon and nitrogen flow through complex anaerobe metabolism. The sealed rotor chamber used in HRMAS NMR maintains an anaerobic environment, ensuring maintenance of reducing conditions, mass conservation, and biological containment. The non-destructive nature of HRMAS NMR spectroscopy enables high-resolution analyses of living cells in small reaction volumes, on the order of tens of microliters, whereas alternatives using GC, MS, or regular solution NMR often require orders of magnitude more volume and cellular mass to support longitudinal sampling and analyses. The acquisition of HRMAS NMR datasets is also far more rapid and does not require further extraction or preparation beyond loading of living cells in the rotor.

Longitudinal tracking of fermentable ^13^C-labeled substrates supports high-resolution monitoring of individual pathways within complex nutrient conditions. The cellular scale of genome-scale metabolic models provides means to incorporate multiple NMR datasets tracking fermentable substrates into a unified dFBA solution with inference of flux through pathways not directly captured by NMR analyses, and of cellular or experimental objectives that may include ATP generation, biomass, or production of industrially important metabolites. ^13^C-NMR analyses of ^13^C- and ^15^N-labeled substrates quantified ^15^N flow to ^13^C backbones, confirming integration points identified by dFBA. We apply this framework to identify electron- and nitrogen-cycling systems driving ATP production, including promising gene-level targets for downstream therapeutic interventions against *C. difficile* infection.

To overcome a prior limitation in using NMR time series data for genome-scale predictions of cellular metabolism, we present three approaches to transform integrated ^13^C-NMR trajectories into flux estimates to support dFBA simulations. The case of proline provides an example with a known 1:1 correspondence between substrate and its product 5-aminovalerate^21^. In this case, we scaled the 5-aminovalerate signal trajectory to the decrease in the known starting concentration of proline. In the case of leucine, which produces two fermentation metabolites, we scaled the NMR trajectories with GC-derived concentrations of the fermentation products isovalerate and isocaproate in endpoint cultures.

Fermentation of [U-^13^C]glucose provided the most complex case for determining flux estimates from the NMR signals. For one, fermentation of [U-^13^C]glucose proceeds through more complex branching pathways than [U-^13^C]proline or [U-^13^C]leucine. Metabolites produced from glucose fermentation also originate from other metabolic pathways. For example, cellular pools of acetate can include [U-^13^C]acetate from [U-^13^C]glucose and ^12^C acetate from glycine fermentation^38^ and Wood-Ljungdahl acetogenesis^21^. In this complex case involving mixed-source pooling we used ^13^C-NMR of defined standards to estimate the concentrations of ^13^C products by quantifying the molecular context effects on ^13^C signal enhancement. We then calculated estimated endpoint concentrations by scaling the input concentration of the starting fermentation substrate, [U-^13^C]glucose in this case, by the relative integrated signals of the ^13^C products with respect to [U-^13^C]glucose, then correcting by the relative signal enhancement factors determined from the standard solutions. In total, these approaches represent a versatile toolbox that overcomes a critical obstacle to the application of ^13^C-NMR datasets to flux constraints for FBA simulations.

Our use of ^15^N-^13^C J-coupling to amplify less NMR-sensitive ^15^N nuclei with more sensitive ^13^C-NMR presents a conceptual foundation for the simultaneous tracking of atomic species to determine their biological and chemical importance. This concept may be extended to covalent bonding with other NMR-active nuclei such as ^1^H or ^17^O. We leveraged this technique to track two distinct spin-active nuclei in an NMR time series, study interactions between metabolic pathways involving multiple substrates, and experimentally confirm integration points predicted by dFBA.

NMR time series and genome-scale metabolic analyses identified a unique strategy Stickland fermenters employ to integrate Stickland metabolism with high-flux glycolytic metabolism, generating alanine to support cellular growth, energy generation, and more energy-efficient nitrogen handling^13,22^. dFBA simulations predicted a two-phase process for nitrogen flow, the first phase being the release of abundant ammonia from oxidative deamination of amino acids during *Clostridial* Stickland fermentation (Fig. 3d,f,i, Fig. 4, reaction 14, “Nitrogen Cycling”)^31^. The second phase is characterized by the re-assimilation of released ammonia by glutamate dehydrogenase and concomitant transamination of pyruvate to alanine (Fig. 3i, Fig. 4, reactions 13 and 14). HRMAS ^13^C NMR of *C. difficile* cultures grown with [U-^13^C]glucose and L-[^15^N]leucine confirmed predictions of enriched ^15^N flow from leucine to [^13^C,^15^N]alanine formed with ^13^C carbon backbones from [U-^13^C]glucose (Fig. 5c,d, Fig. 4, “Nitrogen cycling”). After the depletion of preferred Stickland acceptors (Fig. 3d,f), alanine’s use as a nitrogen sink consumes reducing equivalents, regenerating oxidized electron carriers for ATP-producing oxidative fermentations (Fig. 3i, Fig. 4, reactions 13 and 14)^22^, and further supports cellular systems in protein and peptidoglycan synthesis^39^, energy storage^21,40^, and osmotic balance^41,42^. Our findings also inform *in vivo* behaviors of *C. difficile* by illustrating how the pathogen balances concurrent glycolytic and amino acid fermentation pathways when presented with abundant fermentable carbohydrates and amino acids, conditions that occur *in vivo* after antibiotic ablation of the microbiota^24^. This analytic approach can support further analyses of prokaryotic physiology including microbial responses to antibiotics, or optimization of conditions to produce industrially important chemicals from different input feedstocks. Living cell HRMAS ^13^C NMR provides an unified methologic approach to define cellular-scale anaerobic metabolism for diverse applications.

## Supporting information

icdf metabolic model updates

## Methods

### Strains

A pathogenicity locus (PaLoc)-deleted strain of *Clostridioides difficile* ATCC 43255 was generated that lacked the *tcdB, tcdE* and *tcdA* genes to reduce biohazard risks for NMR analyses. The deletion mutant was created using a toxin-mediated allele exchange method^43^. Briefly, two regions of approximately 800 bp of DNA flanking the region to be deleted were amplified by PCR from *C. difficile* ATCC43255 using the primers in Supplementary Table 9. Purified PCR products were cloned into the PmeI site of the pMSR0 vector using NEBuilder HiFi DNA Assembly. The resulting plasmid was transformed into E. coli strain NEB10β (New England Biolabs) and the insert verified by sequencing. E. coli strains were cultured aerobically at 37 °C in LB broth or LB agar supplemented with chloramphenicol (15 μg/ml). The plasmid was then transformed into *E. coli* HB101 (RP4) and conjugated into *C. difficile* ATCC43255 that had been heat-shocked at 50°C for 15 min beforehand. Transconjugants were selected on BHI agar plates supplemented with cycloserine (250 μg/ml), cefoxitin (25 μg/ml), and thiamphenicol (15 μg/ml). Allelic exchange was performed as described^43^. This strain was shown to be non-toxigenic *in vitro* to fibroblast cell culture using the methods described in Girinathan *et al*.^24^.

### Strain culture conditions

The ΔPaLoc strain of ATCC 43255 was cultured for 12 hours in supplemented brain-heart infusion media (BHIS; Remel, Lenexa, KS, USA). Cells were spun and washed three times in pre-reduced PBS, prepared in molecular-clean water, and diluted to introduce 100,000 cells into HRMAS NMR rotor inserts for analyses. Preparations were serially diluted and plated to Brucella agar (Remel, Lenexa, KS, USA) to quantitate vegetative cells and spores used in input preparations. Spore counts after 12 hours of culture in BHIS were <0.1% of vegetative cells.

*C. difficile* Modified Minimal Medium (MMM; pH 7.2) supplemented with 100 µM sodium selenite was prepared as described^25^ with [U-^13^C]glucose (D-glucose-^13^C_6_, 99 atom), L-[U-^13^C]proline (L-proline-^13^C_5_, 99 atom %), or L-[U-^13^C]leucine (L-leucine-^13^C_6_, 98 atom %) as indicated in Supplementary Table 1 (Sigma-Aldrich, St. Louis, MO, USA). Studies evaluating flow of amino nitrogen from Stickland-fermented leucine to [U-^13^C]pyruvate generated in glycolysis used L-[^15^N]leucine (Sigma-Aldrich, St. Louis, MO, USA).

NMR rotors were loaded in an anaerobic chamber with approximately 100,000 CFU added to the defined MMM conditions. The insert was sealed and removed from the chamber for NMR analyses.

After analyses, rotor contents were checked for pH, serially diluted, and plated to Brucella agar and incubated anaerobically at 37°C to evaluate vegetative and spore CFU and confirm absence of contaminating species. Cellular growth in the rotor was approximately 30% over the input biomass. The pH remained ranged from 7.17-7.27 after 36h of analyses.

### HRMAS NMR

HRMAS NMR measurements were performed on a Bruker Avance III HD 600 MHz spectrometer (Bruker BioSpin Corporation, Billerica, MA, USA). The sealed Kel-F insert with live cells loaded in the anaerobic chamber was placed with 2 uL of D2O (for field locking) in a 4 mm zirconia rotor, before the rotor was sealed and introduced into the triple-resonance HRMAS probe. One and two dimensional ^1^H and ^13^C NMR were conducted at 37 °C with a spin-rate of 3600 ± 2 Hz. One-dimensional time series spectra were measured alternately and continuously for ^1^H NOESY (Nuclear Overhauser Effect SpectroscopY) with water suppression (∼13 mins) and for proton decoupled ^13^C (∼43 mins) throughout the length of the experimental time. Two-dimensional ^1^H COSY (COrrelated SpectroscopY; ∼3 hrs and 49 mins), proton decoupled ^13^C COSY (∼3 hrs and 30 mins), and ^13^C-decoupled ^1^H-^13^C HSQC (Heteronuclear Signal Quantum Coherence; ∼3 hrs and 38 mins) spectra were inserted in the between the 1D time series. MRS spectra were processed using the TopSpin 3.6.2 (Bruker BioSpin Corporation, Billerica, MA, USA), as well as with NUTS (Acorn NMR Inc., Livermore, CA, USA). All free induction decay files will be made available on the Metabolomics Workbench (https://www.metabolomicsworkbench.org/).

### Identification of metabolite productions with 2D NMR

^13^C labeled metabolites produced from [U-^13^C]glucose, L-[U-^13^C]proline, and L-[U-^13^C]leucine in live cells were identified through 2D NMR (Extended Data Fig. 2) according to reported ^13^C and ^1^H chemical shift values availed from Human Metabolome Database (HMDB, https://hmdb.ca) and Biological Magnetic Resonance Bank (BMRB, https://bmrb.io).

### ^1^H and ^13^C spectra analyses

Individual free induction decay (fid) files were processed using NMRPipe^44^, nmrglue^45^, and in-house python scripts. Fourier transformed spectra were normalized by the noise root-mean-square error of the sparse 130-160 ppm region (Supplementary Figs. 1, 2, and 3). The normalized spectrum stack was rendered as a surface plot in MATLAB R2019b (MathWorks, Natick, MA, USA) with face lightness mapped to the log base-2 of signal. Peaks with height ≥6 times the noise root-mean-square error (130-160 ppm) and separated from other peaks by 0.08 ppm were classified as detectable signals and assigned to compounds using the following algorithm:

1. Subpeaks within 0.3 ppm of each other were clustered.
2. Reference shifts for expected compounds within 0.45 ppm of the cluster were associated with the cluster.
3. All subpeaks from a cluster that associated with a single reference peak were assigned to the compound producing that reference peak.
4. In cases where a single cluster was associated with multiple reference peaks, subpeaks were manually assigned to compounds according to the reference chemical shifts from HMDB and associated splitting patterns. When the number of detected subpeaks was less than the expected multiplicity of the contributing reference peaks, significant overlap was suspected and the clusters were excluded from analyses, as was the case for the isovalerate and isocaproate shifts at ∼24.6 ppm in the [U-^13^C]leucine experiment. The remainder of manually assigned clusters used the 1D ^13^C and 1D ^1^H-^13^C HSQC spectra to deconvolute subpeaks in cases with significant overlap.

Assigned peak signals were concatenated into signal ridges, color-labeled by metabolite, and superimposed over the stack as a scatter plot with stems to the *xy* plane and a smoothing spline curve fit. Surface regions within 0.5 ppm of each reference peak were colorized.

Assigned peaks ≤100 ppm were curve-fit using a Voigt (Gaussian-Lorentzian convolution) lineshape and integrated. Logistic curve fits of metabolite integral vs. time were calculated in python using a least-squares regression (SciPy^46^ 1.6.2) according to equation (1) (Supplementary Table 2). The custom python and MATLAB scripts are available via GitHub (https://github.com/Massachusetts-Host-Microbiome-Center/nmr-cdiff).

### Metabolic modeling

A previously published genome-scale metabolic model of *C. difficile* strain 630, icdf834^30^, was modified using the COBRApy toolbox^47^ and custom python scripts. Exchange reaction bounds were set to 3% of the millimolar concentration of media components used experimentally (Supplementary Table 1). The following changes were made based on experimental data and to support biologically relevant processes energetically and thermodynamically. The updated model, icdf843, is included within the GitHub repository.

1. An exchange reaction was added for iron(II), transport and secretion reactions were added for propionate, phenylacetate, indole-3-acetate, butyrate, and n-butanol, and proton-motive force dependent transport reactions were added for acetate, L-alanine, L-leucine, L-proline, L-isoleucine, isovalerate, isobutyrate, 2-methylbutyrate, isocaproate, and 5-aminovalerate (not listed in Supplementary Table 10).
2. A reaction representing the hydrolysis of ATP was added as an objective function for dFBA analyses. (Supplementary Table 10, row 1).
3. Reactions of the oxidative Stickland pathways were added, employing ferredoxin as the oxidizing agent. An electron-bifurcating reaction for the reduction of isocaprenoyl-CoA was added (Supplementary Table 10, rows 2-4)^19^.
4. A proton motive force (PMF)-generating Rnf complex reaction and an ATP synthase reaction were added (Supplementary Table 10, rows 5-6) to complete the Stickland energy salvage mechanism^21^.
5. Proposed pathways for butyrate and a putative propionate fermentation were added (Supplementary Table 10, rows 7-13) based on experimental evidence of secretion in *C. difficile*^21^.
6. Reactions for formate dehydrogenase and an electron-bifurcating hydrogenase were added with appropriate electron carriers (Supplementary Table 10, rows 14-15)^22^.
7. Reversibility of some existing reactions were altered to reflect practical thermodynamic bounds and prevent cycles from occurring in the flux balance analysis (FBA) solution (Supplementary Table 10, rows 17-35).
8. Reactions were removed per one or more of the following critiera:
  a. Reactions causing thermodynamically infeasible FBA cycles (Supplementary Table 10, rows 35-38)
  b. Reactions superseded by those added in steps 1-7 (Supplementary Table 10, rows 39-45)
  c. Reactions utilizing incorrect oxidizing or reducing agents (Supplementary Table 10, rows 46-49)
  d. Reactions lacking experimental and genomic evidence for their presence in *C. difficile* (Supplementary Table 10, rows 50-end).

### Estimation of exchange fluxes for dFBA

The simplest approach, used in cases where a 1:1 relationship between the labeled substrate and its product eliminates the need for a product flux constraint, estimates substrate exchange flux using a logistic equation scaled by a known input substrate concentration. In *C. difficile*, proline fermentation is well-defined and is only known to follow a single pathway, yielding 5-aminovalerate^18^, making it an ideal candidate for this approach. A minority of proline is also utilized for protein synthesis. Given the known concentration of proline in MMM, we transformed the logistic derivative equation (3) for proline into a flux trajectory by multiplying it by the factor [Proline]_*init*_/(*L*_Proline_ + *C*_Proline_), where [Proline]_*init*_ is the initial proline concentration in the NMR run and *L*_Proline_ and *C*_Proline_ are logistic equation (1) coefficients for proline. We left the 5-aminovalerate flux unconstrained, assuming that the vast majority of proline would flow through this pathway, with minimal flux through the biomass reaction. This approach performs best for compounds like proline that possess a single, well-defined fermentation pathway, removing the need for a constraint on the product’s exchange flux.

In cases where a labeled substrate is metabolized into multiple products, such as leucine’s fermentation via two separate pathways, we estimated exchange fluxes for the products isocaproate and isovalerate by determining an expected ratio using gas chromatography (GC) with flame ionization detection (FID). Volatile short-chain fatty acids were extracted and quantified from stationary phase cultures of *C. difficile* ATCC43255 ΔPaLoc grown in MMM lacking isoleucine as described in Girinathan *et al*.^24^, as shown in Supplementary Table 3. Expected isocaproate and isovalerate yields (*Y*_Product_) per mole of leucine were estimated by taking the molar ratio of each product in the FID readout to the input leucine. Flux trajectories for the NMR runs were then estimated by multiplying equation (3) for each product by the factor [Leucine]_*init*_\**Y*_Product_/*L*_Product_, where [Leucine]_*init*_ is the initial leucine concentration in the NMR run.

In the third example, we estimated exchange fluxes of [U-^13^C]glucose metabolites using the input glucose concentration, the relative ^13^C-NMR peak areas of glucose and each metabolite, and correction factors for NOE-induced signal enhancement empirically derived from standard solutions containing U-^13^C metabolites. Since glucose fermentation is highly integrated with central carbon metabolism, methods to estimate product concentration specific to the ^13^C input substrate are required, such as acetate originating from glucose versus glycine fermentation. For this case we measured standard solutions within the dynamic range of our experiments under ^13^C-NMR and integrated the peak areas to estimate a relationship between concentration and signal amplitude for each compound. As discussed in “Results: Dynamic Flux Balance Analysis”, ^13^C-NMR signal amplitude is dependent on the molecular context of an individual ^13^C atom and is influenced by predictable factors, the most salient being ^1^H-^13^C NOE and spin-rotation^14^. We assume that the relative effects of these properties on ^13^C-NMR signal amplitude remain consistent between ^13^C atoms regardless of NMR acquisition parameters, and thus we define ^*13*^*C-NMR signal enhancement* as the relative NMR signal-concentration ratio of each product with respect to [U-^13^C]glucose. Solutions listed in Supplementary Table 4 were measured by ^13^C-NMR and processed using the procedures in “HRMAS NMR” and “^1^H and ^13^C spectra analyses” above. Concentration-to-signal ratios of [U-^13^C]acetate, [U-^13^C]alanine, [U-^13^C]ethanol, and [U-^13^C]butyrate (Sigma-Aldrich, St. Louis, MO, USA) in each standard solution were plotted vs. the concentration-to-signal ratio of [U-^13^C]glucose in the same standard solution, then fit using a least-squares regression to the equation

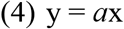

where 1/*a* represents the ^*13*^*C-NMR signal enhancement* of each metabolite with respect to glucose (Extended Data Fig. 1). When low-intensity peaks were excluded (peak area < 18 normalized signal units per ppm), a strong correlation was observed for each compound (R^2^ values displayed on Extended Data Fig. 1), suggesting that the ^*13*^*C-NMR signal enhancement* of each metabolite with respect to glucose is consistent across experiments, even under inconsistent acquisition parameters. A normalization scheme for each product was then derived as follows:

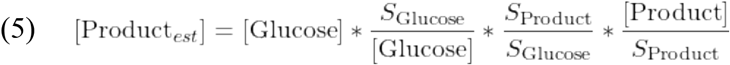

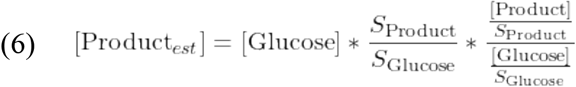

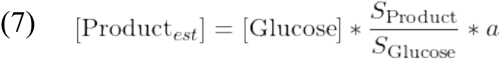

where [Product]_*est*_ is the estimated concentration of the product; [Glucose] is the known input concentration of [U-^13^C]glucose for the run; S_Product_/S_Glucose_ is the ratio of maximum integrated signal between the product and glucose, determined by *f*_Product_(36)/*f*_Glucose_(0) where *f* is the logistic equation (1); and *a* is the product-specific coefficient in equation (4). The logistic derivative equation (3) for each product was then multiplied by [Product]_*est*_/*L*_Product_ and the logistic derivative equation (3) for glucose was multiplied by [Glucose]/(*L*_Glucose_ + *C*_Glucose_) to transform the equations into a flux trajectories. This approach is most suitable for compounds with multiple products where labeled products are commercially available or easily extracted.

### Dynamic Flux Balance Analysis

dFBA was implemented by computing steady-state flux balance analysis (FBA) solutions with time-dependent exchange fluxes using the COBRApy toolbox^47^ and custom python scripts. The rate of isocaproate formation, as represented by the 0.864 ppm peak in the ^1^H-NMR time series, was selected as a normalization marker to align the time axes of all three runs because the 0.864 ppm isocaproate peak is well-separated in the ^1^H-NMR spectra, permitting reliable integration. A logistic curve (1) was fit to the 0.864 ppm isocaproate peak in the ^1^H time series of each run as described in “^1^H and ^13^C spectra analyses.” The time axes for the three runs were normalized by aligning the times at which 5% and 95% of maximum logistic isocaproate ^1^H signal was measured (Extended Data Fig. 5). Exchange (for input metabolites) and secretion (for end metabolites) upper and lower bounds were set to the logistic function derivatives (3) transformed as explained in “Estimation of exchange fluxes for dFBA”, evaluated per time point (Figure 2b,d,f; Supplementary Table 5). Leucine exchange flux was left unconstrained due to low model tolerances. Secretion flux upper bounds of acetate, ethanol, and L-alanine were left unbounded because those products were expected to be non-exclusive to glucose fermentation. As oxidative valine and leucine fermentations co-occur in minimal media^26^, the secretion flux of isobutyrate were locked to the isovalerate trajectory and scaled by the GC-determined isobutyrate/isovalerate ratio. Additionally, acetyl-CoA synthase flux was limited prior to 9 hours, according to *in vitro* results from Gencic et al.^22^. ATP hydrolysis was selected for the biological objective as a common metabolic driver during log phase, stationary phase, and sporulation. Steady-state FBA solutions were calculated along a simulated 36-hour timescale with a resolution of 1 solution per hour. Metabolic flux trajectories of selected reactions were visualized using custom MATLAB scripts (Figure 3). Fluxes contributing to the production and consumption of cytosolic L-alanine, L-glutamate, ATP, pyruvate, and ammonia were recorded at each timepoint (Supplementary Table 6, columns C-AK).

Predicted amino nitrogen flow from L-leucine to L-alanine was estimated by the following scheme. We define two phases of ammonia metabolism: the deamination phase from 1-9 hours where there is net secretion of ammonia, and the assimilation phase from 10-21 hours where there is net uptake of ammonia. To accurately capture the L-alanine amino group contribution from L-leucine, we considered the leucine-origin ammonia released in the deamination phase and recycled in the assimilation phase. For detailed calculations of leucine-origin amino nitrogen flow to alanine, see Supplementary Note 1 and Supplementary Table 6.

### Analyses of ^15^N-^13^C peak splitting in ^13^C NMR spectral analyses of ^13^C alanine

NMR data measured with Bruker BioSpin (Bruker Corporation, Billerica, MA, USA) were analyzed with NUTS, 2D Pro (Acorn NMR Inc., Livermore, CA, USA). The fid files were processed using 0.2Hz line-broadening and were zero-filled before Fourier transformation. After the Fourier transformation, resonance peaks were quantified using curve-fitting with Lorentzian line shapes. Curve fitting was first performed globally to determine the average peak width across all fitted peaks. The curve was then refitted using the width average and Lorentzian function (Extended Data Fig. 3).

### ^13^C NMR analyses of alanine’s α-carbon in media with [U-^13^C]glucose and [^15^N]leucine

The complex peaks from alanine’s α-carbon in the ^13^C NMR spectral region can be deconvoluted based on *(1)* possible ^13^C label patterns (Extended Data Fig. 4A), where *(a)* ^13^C can appear on all three carbons in alanine that results in quadruple peaks occurring at the α-carbon. *(b)* Carbon-13 can also appear on only two carbons in alanine: [1,2-^13^C]alanine and [2,3-^13^C]alanine, while [1,3-^13^C]alanine produces no significant α-carbon 13C NMR signal, and both two ^13^C labeled isotopologues of alanine produce doublet peaks for the α-carbon. *(c)* Carbon-13 only appears on the α-carbon as [2-^13^C]alanine to produce a singlet peak. *(2)* J-coupling of ^15^N-^13^C splits the α-carbon ^13^C NMR peak into two peaks when ^15^N is bound to α-^13^C (Extended Data Fig. 4B). *(3)* the isotope effect that introduces small shifts in the measured resonances^**36**^ (Extended Data Fig. 4D).

## Acknowledgments

We thank Mary Delaney for microbiology support, Michael Judge and Art Edison for helpful comments and for confirming capacity of sealed HRMAS NMR rotors to maintain anaerobic conditions.

## Funding

National Institutes of Health grant R01AI153653 (LB)

National Institutes of Health grant P30DK056338 (LB)

BWH Precision Medicine Institute (LB)

National Institutes of Health grant S10OD023406 (LLC)

National Institutes of Health grant R21CA243255 (LLC)

National Institutes of Health grant R01AG070257 (LLC)

MGH A. A. Martinos Center for Biomedical Imaging (LLC, LB)

## Author contributions

Conceptualization: LB, LLC

Methodology: AP, BG, JP, BD, LLC, LB

Software: AP

Investigation: AP, BG, JP, PAS, BD, IHM, CM, LLC, LB

Formal analysis: AP, IHM, CM, LLC, LB

Funding acquisition: LB, LLC

Writing – original draft: AP, LB

Writing – review & editing: AP, JP, PAS, BD, IHM, CM, LLC, LB

Visualization: AP, IHM, CM, LLC

## Data and materials availability

Primary datasets are available in the supplementary materials. All code is available under the Apache License v2.0 on GitHub at https://github.com/Massachusetts-Host-Microbiome-Center/nmr-cdiff

