## Supplemental Methods and Data for "Elucidating dynamic anaerobe metabolism with Live Cell HRMAS ^13^C NMR and genome-scale metabolic modeling"

**Supplementary Table 1:** ***C. difficile*** **modified minimal media (MMM), adapted from Karasawa et al.**[23] *In conditions containing [U-$^{13}$C]glucose, [U-$^{13}$C]leucine, and/or [$^{15}$N]leucine, the labeled isotopologue was substituted for the natural abundance compound at the same millimolar concentration. **In conditions containing [U-$^{13}$C]proline, [U-$^{13}$C]proline was substituted for natural abundance proline at 30 mM.

| **Stock solution Component** | **Final concentration in MMM** (mg/ml unless indicated) | **Final concentration in MMM** (mM unless indicated) |
|---|---|---|
| D-glucose* | 4.95 | 27.5 |
| L-tryptophan (W) | 0.1 | 0.49 |
| L-isoleucine (I) | 0.3 | 2.29 |
| L-leucine (L)* | 1 | 7.63 |
| L-valine (V) | 0.3 | 2.56 |
| L-cysteine(C) | 0.5 | 4.13 |
| L-proline (P)** | 0.8 | 6.96 |
| L-methionine (M) | 0.2 | 1.34 |
| L-histidine (H) | 0.1 | 0.65 |
| L-arginine (R) | 0.2 | 1.15 |
| Glycine (G) | 0.1 | 1.33 |
| L-threonine (T) | 0.2 | 1.68 |
| $Na_2HPO_4$ | 5 | 35.2 |
| $NaHCO_3$ | 5 | 59.5 |
| $KH_2PO_4$ | 0.9 | 6.6 |
| NaCl | 0.9 | 15.4 |
| $(NH_4)_2SO_4$ | 0.04 | 0.303 |
| $CaCl_2$•$2H_20$ | 0.026 | 0.177 |
| $MgCl_2$•$6H_20$ | 0.02 | 0.099 |
| $MnCl_2$•$4H_20$ | 0.01 | 0.051 |
| $CoCl_2$•$6H_20$ | 0.001 | 0.004 |
| D-biotin | 0.001 | 0.004 |
| Calcium-D-pantothenate | 0.001 | 0.004 |
| $FeSO_4$•$7H_2O$ | 0.004 | 0.014 |
| $Na_2SeO_3$ | 17.3 ng/ml | 100 µM |
| $D_2O$ | 10% v/v | |

**Supplementary Table 2:** Logistic fit coefficients with standard error.

| Metabolite | *L* | *k* | *x0* | *C* |
|---|---|---|---|---|
| **Proline** | 560.205 ± 10.495 | -0.334 ± 0.016 | 11.772 ± 0.145 | 394.989 ± 3.110 |
| 5-aminovalerate | 518.759 ± 3.090 | 0.398 ± 0.019 | 11.993 ± 0.125 | — |
| **Leucine** | 267.553 ± 6.030 | -1.019 ± 0.095 | 6.643 ± 0.106 | -1.829 ± 2.439 |
| Isovalerate | 14.219 ± 0.423 | 1.932 ± 0.766 | 4.590 ± 0.241 | — |
| Isocaproate | 75.773 ± 0.634 | 1.963 ± 0.208 | 7.699 ± 0.062 | — |
| **Glucose** | 1407.383 ± 62.666 | -0.187 ± 0.016 | 16.163 ± 0.415 | 191.084 ± 29.225 |
| Acetate | 773.609 ± 11.678 | 0.211 ± 0.019 | 16.240 ± 0.496 | — |
| Alanine | 334.628 ± 6.416 | 0.309 ± 0.046 | 14.464 ± 0.555 | — |
| Ethanol | 797.009 ± 10.735 | 0.190 ± 0.013 | 22.008 ± 0.422 | — |
| Butyrate | 136.042 ± 3.073 | 0.258 ± 0.033 | 28.571 ± 0.573 | — |

**Supplementary Table 3: Concentrations of isovalerate, isocaproate, and isobutyrate determined by gas chromatography with flame ionization detection (GC-FID).**

| | Volatile acids | | | Proportions | | | |
|---|---|---|---|---|---|---|---|
| **Condition** | **Isovalerate (iVal)** | **Isocaproate (iCap)** | **Isobutyrate (iBut)** | $\frac{\text{iCap}}{\text{iVal}}$ | $\frac{\text{iCap}}{\text{iCap + iVal}}$ | $\frac{\text{iVal}}{\text{iCap + iVal}}$ | $\frac{\text{iBut}}{\text{iVal}}$ |
| A | 2.956 | 5.069 | 1.383 | 1.715 | 0.632 | 0.368 | 0.468 |
| B | 2.899 | 4.899 | 1.361 | 1.690 | 0.628 | 0.372 | 0.469 |
| C | 2.973 | 5.218 | 1.357 | 1.755 | 0.637 | 0.363 | 0.456 |
| D | 3.146 | 5.485 | 1.447 | 1.743 | 0.635 | 0.365 | 0.460 |
| **Average** | **2.994** | **5.168** | **1.387** | **1.726** | **0.633** | **0.367** | **0.463** |

**Supplementary Table 4: Composition of standard solutions containing [U-$^{13}$C]glucose and selected metabolites at defined concentrations.**

| Solution | Concentration (mM) | | | | |
|---|---|---|---|---|---|
| | Glucose | Ethanol | Acetate | Alanine | Butyrate |
| A | 26.85 | 0.00 | 0.00 | 0.00 | 0.00 |
| B | 19.35 | 0.00 | 6.43 | 6.52 | 0.00 |
| C | 11.85 | 2.39 | 14.07 | 12.72 | 2.72 |
| D | 5.87 | 5.22 | 17.96 | 14.78 | 5.45 |
| E | 2.72 | 6.85 | 18.85 | 15.22 | 7.49 |
| F | 10.00 | 10.00 | 13.77 | 10.00 | 12.52 |

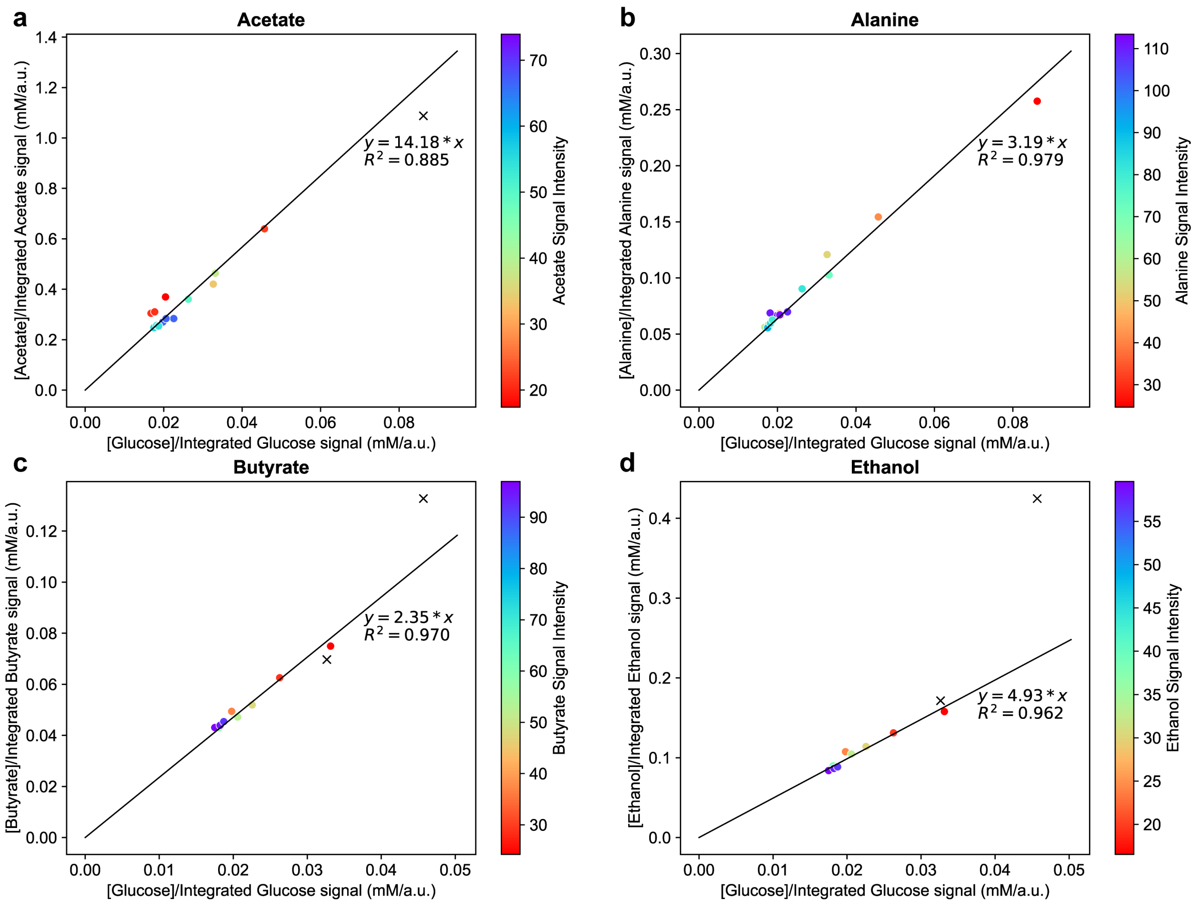


**Extended Data Figure 1: $^{13}$C signal enhancement standard curves of U-$^{13}$C metabolites with respect to glucose.** (**a**) Scatter plot of concentration-signal ratio of [U-$^{13}$C]acetate vs. [U-$^{13}$C]glucose, determined from HRMAS $^{13}$C-NMR acquisitions of the solutions listed in Supplementary Table 4. Only peaks ≤ 100 ppm are included in the analyses; acquisitions where the average per-carbon peak area for a compound is less than 18 are marked with a black cross and excluded from the linear regression. Best fit line shown with equation and R-squared statistic. (**b**) Scatter plot of concentration-signal ratio of [U-$^{13}$C]alanine vs. [U-$^{13}$C]glucose, determined as in panel (a). (**c**) Scatter plot of concentration-signal ratio of [U-$^{13}$C]butyrate vs. [U-$^{13}$C]glucose, determined as in panel (a). (**d**) Scatter plot of concentration-signal ratio of [U-$^{13}$C]ethanol vs. [U-$^{13}$C]glucose, determined as in panel (a).

**Supplementary Table 5: Logistic coefficients for the estimated concentration curves constraining dFBA analyses.**

| Metabolite | *L* | *k* | *x0* | *C* |
|---|---|---|---|---|
| **Proline** | 4.082 ± 0.076 | -0.334 ± 0.016 | 11.772 ± 0.145 | 2.878 ± 0.023 |
| 5-aminovalerate | 3.780 ± 0.023 | 0.398 ± 0.019 | 11.993 ± 0.125 | — |
| **Leucine** | 7.683 ± 0.173 | -1.019 ± 0.095 | 6.643 ± 0.106 | -0.053 ± 0.070 |
| Isovalerate | 2.800 ± 0.083 | 1.932 ± 0.766 | 4.590 ± 0.241 | — |
| Isobutyrate | 1.297 ± 0.039 | 1.932 ± 0.766 | 4.590 ± 0.241 | — |
| Valine | 1.297 ± 0.039 | -1.932 ± 0.766 | 4.590 ± 0.241 | — |
| Isocaproate | 4.830 ± 0.040 | 1.963 ± 0.208 | 7.699 ± 0.062 | — |
| **Glucose** | 24.459 ± 1.089 | -0.187 ± 0.016 | 16.163 ± 0.415 | 3.321 ± 0.508 |
| Acetate | 13.445 ± 0.203 | 0.211 ± 0.019 | 16.240 ± 0.496 | — |
| Alanine | 5.816 ± 0.112 | 0.309 ± 0.046 | 14.464 ± 0.555 | — |
| Ethanol | 13.851 ± 0.187 | 0.190 ± 0.013 | 22.008 ± 0.422 | — |
| Butyrate | 2.364 ± 0.053 | 0.258 ± 0.033 | 28.571 ± 0.573 | — |

**Supplementary Table 6: Selected dFBA-predicted metabolic fluxes.** Predicted producing and consuming fluxes for L-alanine (alaL_c, producing: columns C-D, consuming: column E), L-glutamate (gluL_c, producing: columns F-H, consuming: columns I-J), ATP (atp_c: columns K-S, consuming: columns T-AA), pyruvate (pyr_c: columns AB-AC, consuming: columns AD-AF), and ammonia (nh3_c: columns AG-AK, consuming: columns AK-AL). The model objective flux is in column U. Calculations for estimated amino group flux from leucine to alanine are in columns AM-BB.

**Supplementary Note 1:** To accurately capture the L-alanine amino group contribution from L-leucine, we considered the leucine-origin ammonia released in the deamination phase and recycled in the assimilation phase. First, the leucine-origin ammonia influx during the deamination phase (Supplementary Table 6, column AT) was defined at each timepoint as the proportion of L-glutamate amino nitrogen from L-leucine (Supplementary Table 6, column AN) times the proportion of ammonia from glutamate (Supplementary Table 6, column AQ) times the outflux of ammonia (Supplementary Table 6, column AS). The proportion of leucine-origin pooled ammonia at the beginning of the assimilation phase (Supplementary Table 6, cell AS45) was calculated as the cumulative sum of the ammonia pooled from L-leucine (Supplementary Table 6, cell AS42) divided by the total ammonia pooled after 9 hours (Supplementary Table 6, cell AS44).

At each timepoint during the assimilation phase, the leucine-origin glutamate flux was assumed to have two components; first, the amino groups transaminated directly from leucine at the timepoint (nonzero only before 13 hours), and secondly, the re-assimilation onto glutamate of leucine-origin ammonia released during the deamination phase. The first pool is equal to the flux through the branched-chain amino acid transaminase (Supplementary Table 6, column AL), and the second pool is equal to the product of the leucine-origin ammonia proportion (Supplementary Table 6, cell AS45) and the glutamate assimilation flux from ammonia (Supplementary Table 6, column AT). The proportion of leucine-origin glutamate (Supplementary Table 6, column AW) was taken as the sum of these two factors (Supplementary Table 6, column AV) divided by the total glutamate influx (Supplementary Table 6, column AM).

Lastly, the amino group flux from leucine to alanine (Supplementary Table 6, column AY) was defined as the leucine-origin glutamate proportion (Supplementary Table 6, column AW) times the amino group flux from glutamate to alanine via ALT (Supplementary Table 6, column AX). The total proportion of alanine with leucine-origin amino groups (Supplementary Table 6, cell AZ43) was estimated as the cumulative amino group flux from leucine to alanine (Supplementary Table 6, cell AY41) as a proportion of total alanine synthesis (Supplementary Table 6, cell AZ41).

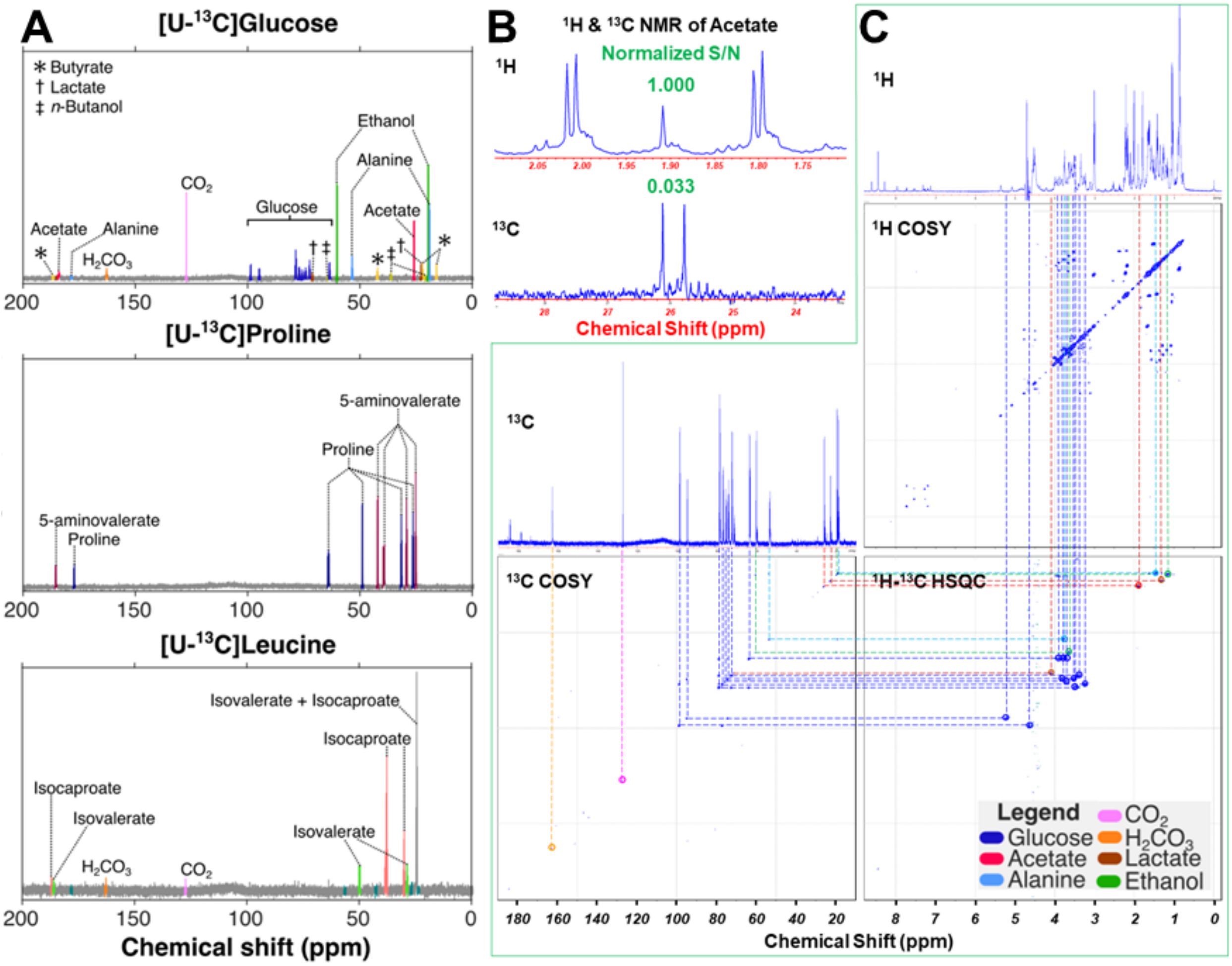


**Extended Data Figure 2: Live cell $^{13}$C NMR spectra and metabolite identification**. (**A**) Proton-decoupled $^{13}$C NMR spectra of live cells following reactions with [U-$^{13}$C]glucose, L-[U-$^{13}$C]proline, and L-[U-$^{13}$C]leucine, respectively. (**B**) $^{1}$H and proton-decoupled $^{13}$C NMR spectra of [U-$^{13}$C]glucose produced $^{13}C_2$-acetate, measured at 45.5 and 45.0 hours, respectively. Relative signal-to-noise ratio (S/N) normalized to unit time for measured for $^{1}$H (S/N = 1.000) and $^{13}$C (S/N = 0.033). The double-doublet peaks seen in the $^{1}$H spectrum resulted from $^{1}$H J-couplings with both methyl and carboxyl $^{13}$C nuclei. The center peak represented methyl-$^{1}$H bond to $^{12}$C. The doublet peaks seen in the $^{13}$C spectrum were the result of $^{13}$C-$^{13}$C J-coupling between methyl and carboxyl $^{13}$C nuclei. (**C**) An illustration of metabolite identifications for live cells grown with [U-$^{13}$C]glucose through 2D $^{13}$C COSY, $^{13}$C-$^{1}$H HSQC, and $^{1}$H COSY according to reported $^{13}$C and $^{1}$H chemical shift values from the Human Metabolome Database (HMDB). In this illustration, the $^{1}$H 1D spectrum was measured at 45.5 hours; $^{1}$H 2D COSY, at 32.6 hours; $^{1}$H-$^{13}$C HSQC, at 27.9 hours; $^{13}$C COSY, at 37.2 hours; and $^{13}$C 1D spectrum was a composite spectrum of all proton-decoupled $^{13}$C spectra (n=11) measured between 24.8 and 45.0 hours.

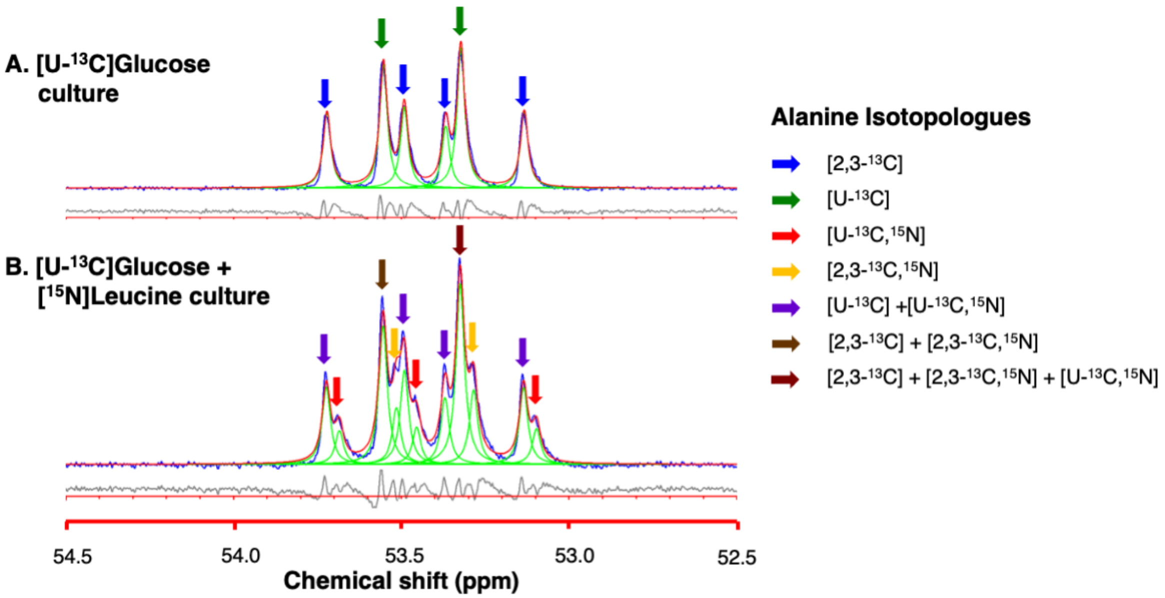


**Extended Data Figure 3: Curve-fitted $^{13}C$ NMR spectral region of alanine's α-carbon following the growth of live *C. difficile* in [U-$^{13}$C]glucose with natural abundance leucine or [$^{15}$N]leucine.** (**A**) Spectrum of culture grown in media containing natural abundance leucine. (**B**) Spectrum of culture grown in media containing [$^{15}$N]leucine. In each resonance spectrum, blue indicates the curve of the experimental raw data; red indicates the overall fitted curve; and green indicates the individual fitted peaks; the grey line indicates the difference between the experimental and fitted curves.

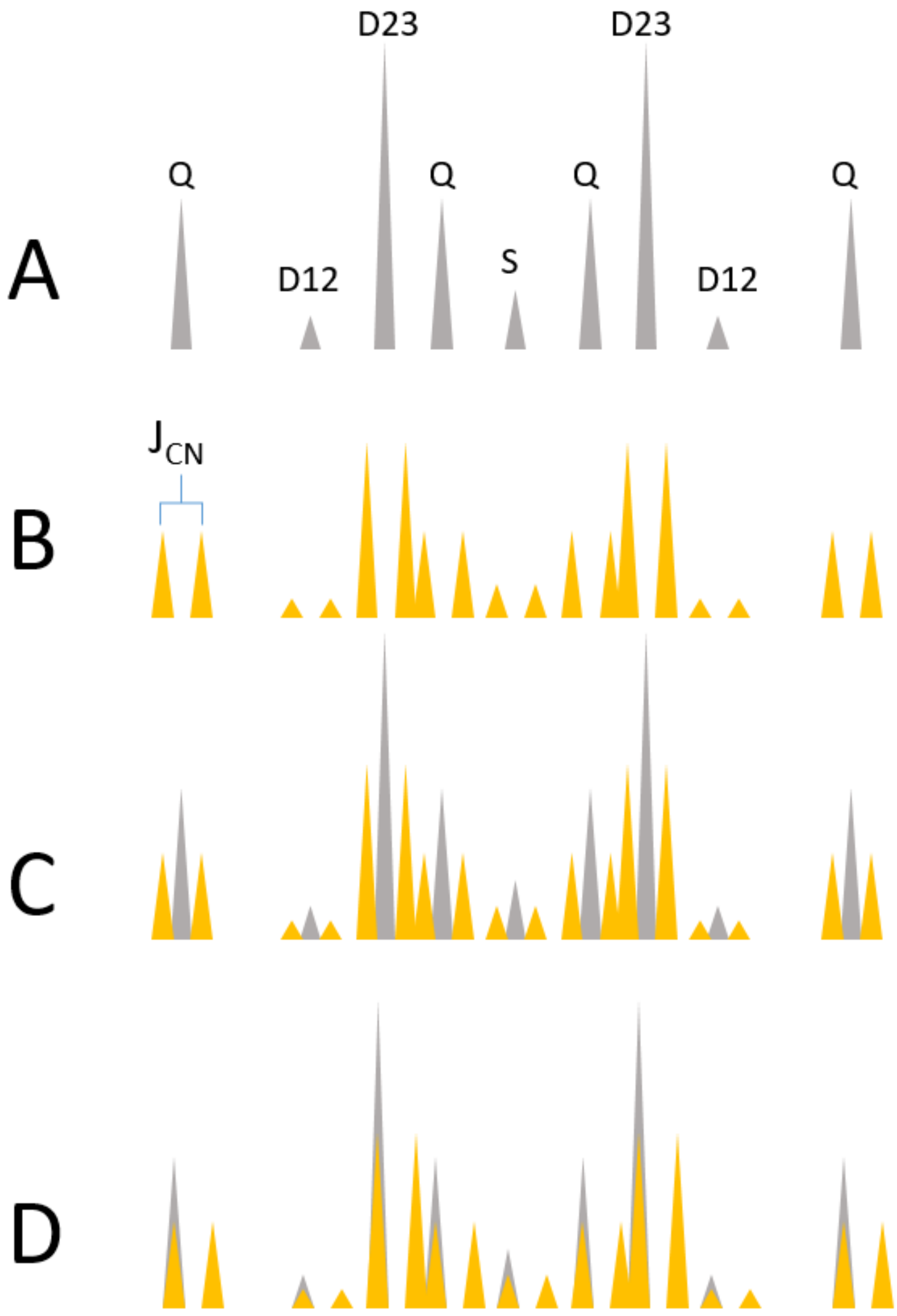


**Extended Data Figure 4: Deconvolution of alanine's α-carbon in the $^{13}C$ NMR spectral region.** (**A**) All possible $^{13}C$ patterns of alanine's α-carbon, as produced from [U-$^{13}C$]glucose, are shown in gray (Q: quadruple peaks from [U-$^{13}C$]alanine, D12 and D23: doublets from [1,2-$^{13}C$]alanine and [2,3-$^{13}C$]alanine, respectively, S: singlet from [2-$^{13}C$]alanine. (**B**) All peaks split into two equal intensity peaks due to $^{15}N$-$^{13}C$ J-coupling, shown in yellow. (**C**) The sum of peaks in A and B, (**D**) Predicted peak profile of all possible alanine species, with calculations that include shift of $^{15}N$ associated $^{13}C$ resonance due to isotope effects.

**Supplementary Table 7. Isotopologue proportions of alanine in stationary phase *C. difficile* cultures grown in media containing [U-$^{13}$C]glucose and either natural abundance leucine or [$^{15}$N]leucine.**

| Alanine | Run1 | Run2 | Run3 | AVG | STDEV | SEM |
|---|---|---|---|---|---|---|
| **U-$^{13}$C, $^{14}$N** | 19% | 24% | 23% | 22% | 2.65% | 1.53% |
| **2,3-$^{13}$C, $^{14}$N** | 17% | 22% | 24% | 21% | 3.61% | 2.08% |
| **U-$^{13}$C, $^{15}$N** | 36% | 27% | 27% | 30% | 5.20% | 3.00% |
| **2,3-$^{13}$C, $^{15}$N** | 28% | 26% | 26% | 27% | 1.15% | 0.67% |
| **TOTAL $^{14}$N** | 36% | 46% | 47% | 43% | 6.08% | 3.51% |
| **TOTAL $^{15}$N** | 64% | 53% | 53% | 57% | 6.35% | 3.67% |

**Supplementary Table 8: Pathways and enzyme systems involved in the metabolism of glucose, leucine, and proline in *Clostridioides difficile*.** Column labels indicate the following information. Number: corresponding reaction numbers shown in Figures 2 and 3. System or Enzyme: corresponding cellular system or enzyme-catalyzed pathway. Gene-association: CD630: associated geneID in the *C. difficile* reference strain CD630. Gene-association: ATCC43255: associated geneID in the *C. difficile* strain ATCC43255 used in the present studies (see Excel).

**Supplementary Table 9: Primers used for construction of the PaLoc- strain.**

| Primer | Sequence (5’ to 3’)* | Use |
|---|---|---|
| BD013 | ttttttgttaccctaagtttGGATGATTTTATGCAAAAGTC | 5’ left arm for *tcdBEA* deletion |
| BD014 | tattttagccCATAAAATTTTCTCCTTTACTATAATATTTTTAC | 3’ left arm for *tcdBEA* deletion |
| BD015 | aaattttatgGGCTAAAATATATGTTTGATAAAAAATTATTC | 5’ right arm for *tcdBEA* deletion |
| BD016 | agattatcaaaaaggagtttCCAGCTTGTTCTGAAGAC | 3’ right arm for *tcdBEA* deletion |
| BD017 | GGAGGATATATAAAAGAGTTTTAGC | 5’ screening of *tcdBEA* deletion |
| BD018 | GGGTATTGCTCTACTGGC | 3’ screening of *tcdBEA* deletion |

*Lowercase bases indicate overlapping sequences

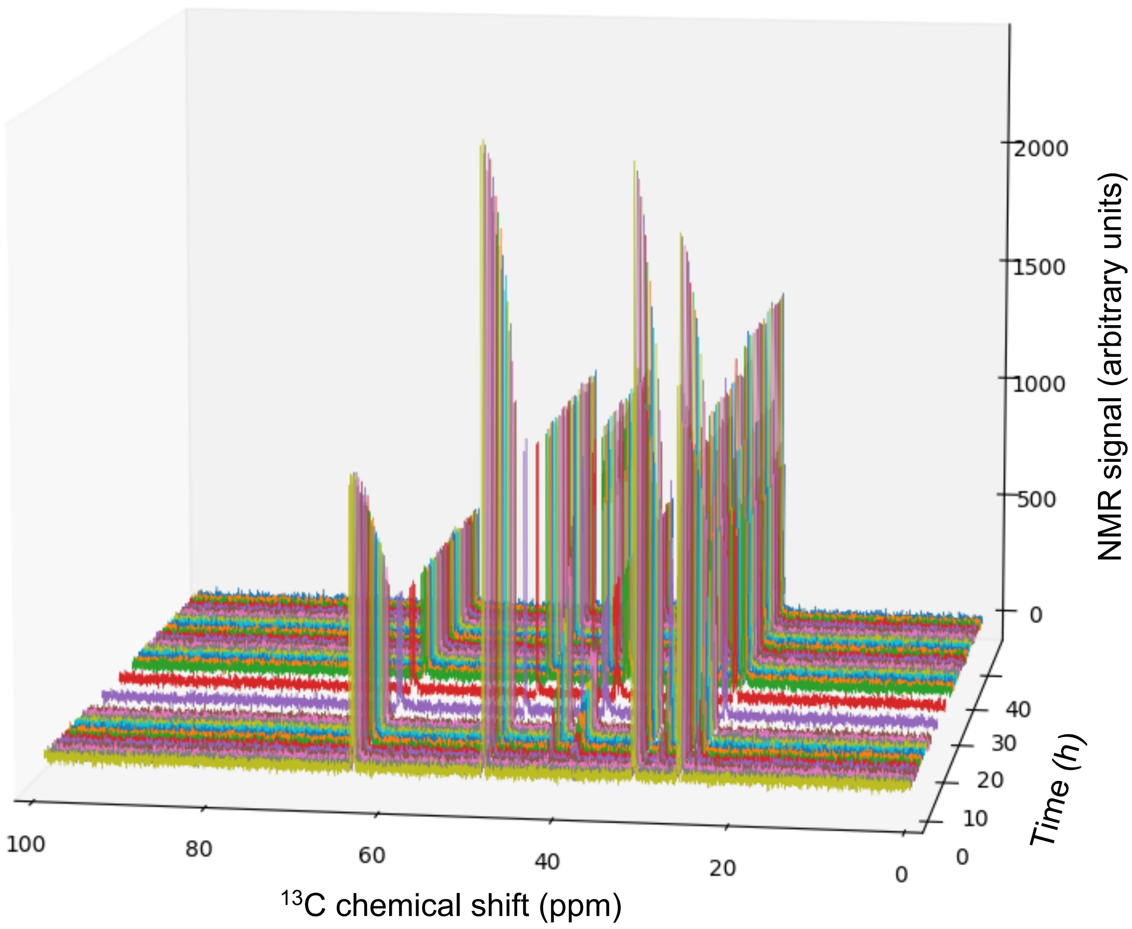


**Supplementary Figure 1: Processed $^{13}$C-NMR spectra from the metabolism of [U-$^{13}$C]proline.**

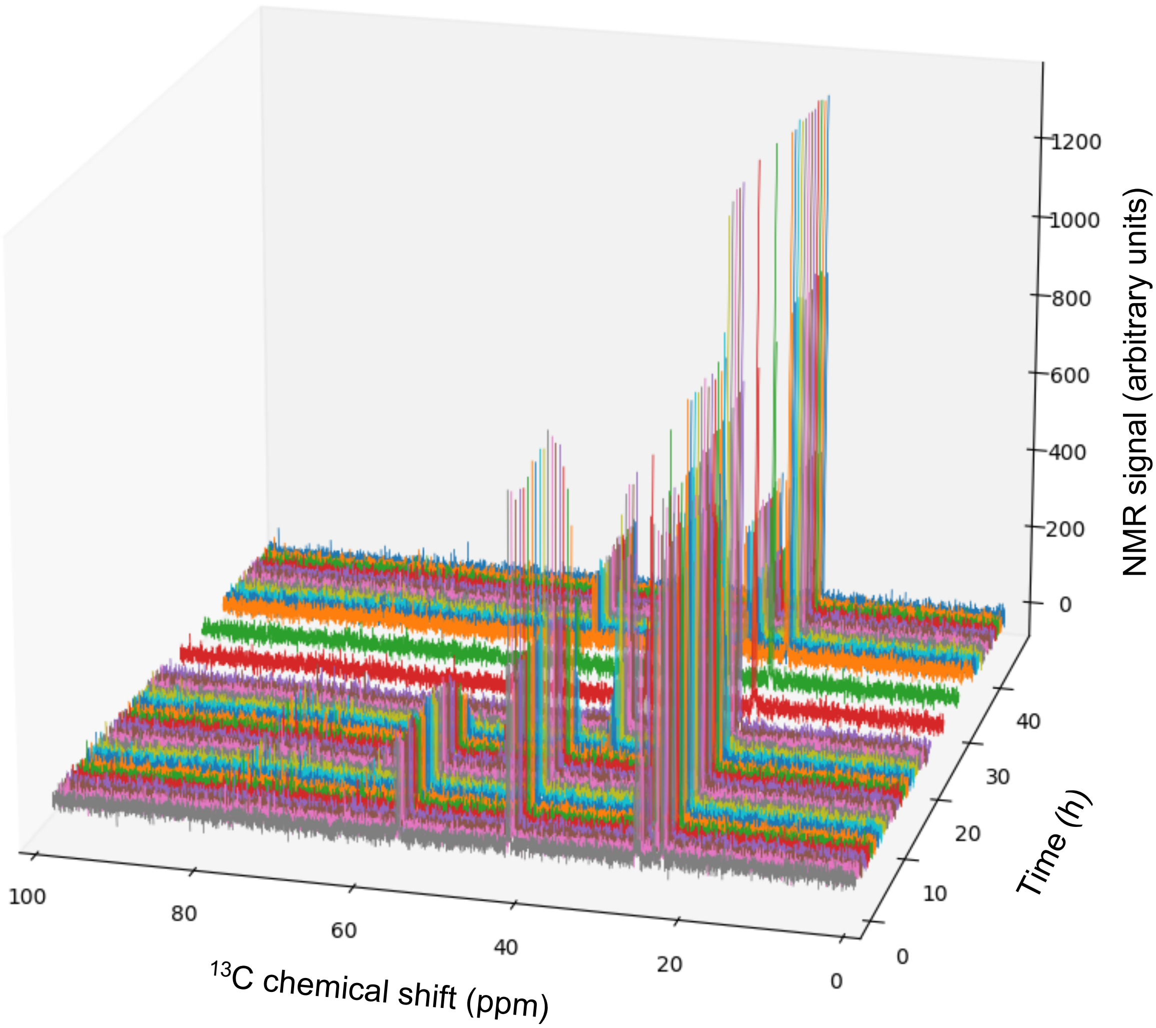


**Supplementary Figure 2: Processed $^{13}$C-NMR spectra from the metabolism of [U-$^{13}$C]leucine.**

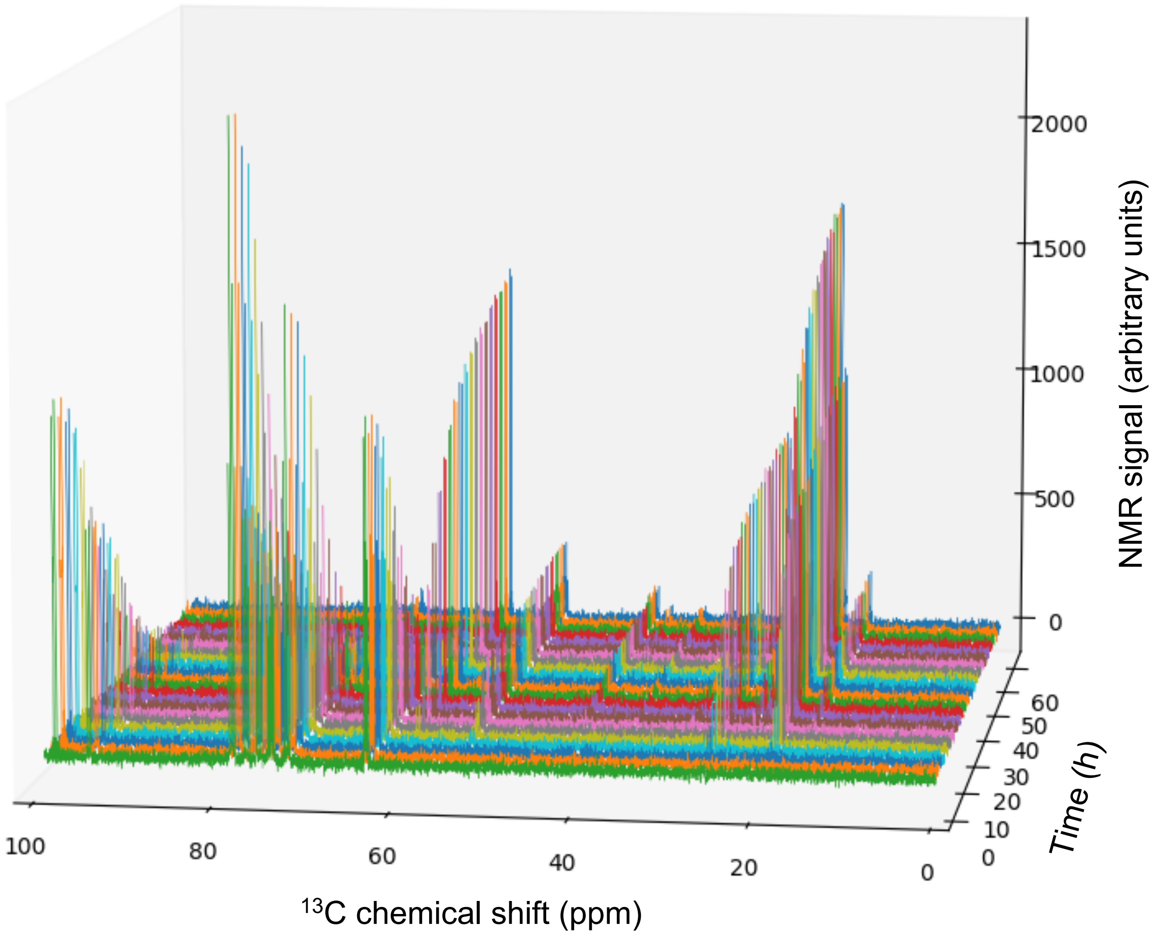


**Supplementary Figure 3: Processed $^{13}$C-NMR spectra from the metabolism of [U-$^{13}$C]glucose.**

**Supplementary Table 10:** Modifications to icdf834, a previously published model of *C. difficile*[28]. The resulting model is named icdf843. Modifications are further elaborated in the Methods (see Excel).

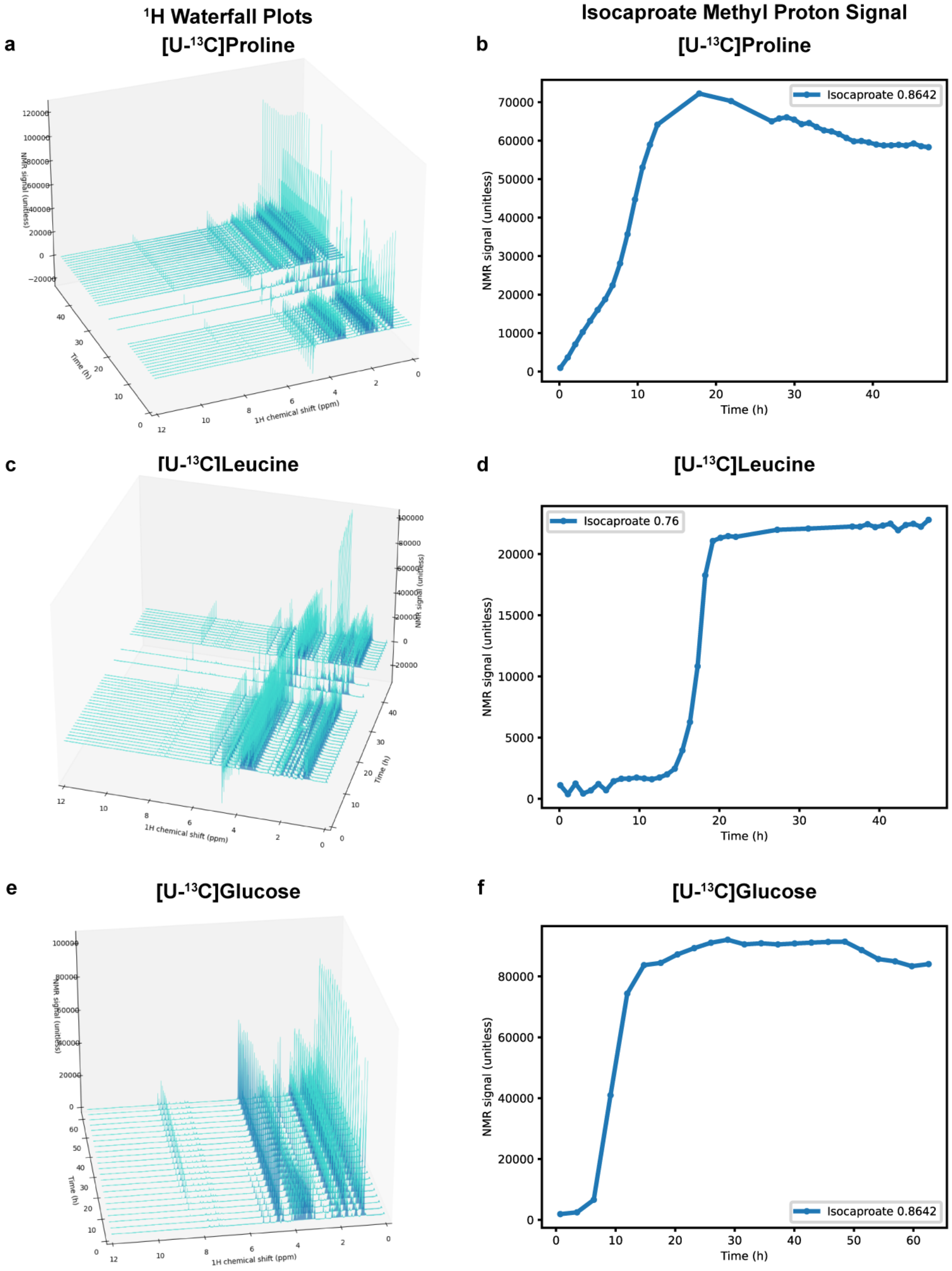

a
$^1$H Waterfall Plots
[U-$^{13}$C]Proline
NMR signal (unitless)
Time (h)
1H chemical shift (ppm)
b
Isocaproate Methyl Proton Signal
[U-$^{13}$C]Proline
Isocaproate 0.8642
NMR signal (unitless)
Time (h)
c
[U-$^{13}$C]Leucine
NMR signal (unitless)
Time (h)
1H chemical shift (ppm)
d
[U-$^{13}$C]Leucine
Isocaproate 0.76
NMR signal (unitless)
Time (h)
e
[U-$^{13}$C]Glucose
NMR signal (unitless)
Time (h)
1H chemical shift (ppm)
f
[U-$^{13}$C]Glucose
Isocaproate 0.8642
NMR signal (unitless)
Time (h)

**Extended Data Figure 5: Processed 1H spectra and isocaproate trajectories.** HRMAS $^{1}$H-NMR waterfall plot of growth in MMM with 30mM [U-$^{13}$C]proline. X-axis shows $^{1}$H NMR chemical shift (ppm), Y-axis: time (hours), and Z-axis: NMR signal (unitless). (**b**) Line plot depicting signal of the isocaproate methyl proton peak vs. time in the [U-$^{13}$C]proline experiment. (**c**) HRMAS $^{1}$H-NMR waterfall plot of growth in MMM with 7.6mM [U-$^{13}$C]leucine; axes as in panel A. (**d**) Line plot depicting signal of the isocaproate methyl proton peak vs. time in the [U-$^{13}$C]leucine experiment. (**e**) HRMAS $^{1}$H-NMR waterfall plot of growth in MMM with 27.5mM [U-$^{13}$C]glucose; axes as in panel A. (**f**) Line plot depicting signal of the isocaproate methyl proton peak vs. time in the [U-$^{13}$C]glucose experiment.
